# Mapping Complex Brain Torque Components and Their Genetic and Phenomic Architecture in 24,112 healthy individuals

**DOI:** 10.1101/2021.03.09.434625

**Authors:** Lu Zhao, William Matloff, Yonggang Shi, Ryan P. Cabeen, Arthur W. Toga

## Abstract

The mechanisms determining the development and individual variability of brain torque (BT) remain unclear. Here, all relevant components of BT were analyzed using neuroimaging data of up to 24,112 individuals from 6 cohorts. Our large-scale data confirmed the population-level predominance of the typical anticlockwise torque and suggested a “first attenuating, then enlarging” dynamic across the lifespan primarily for frontal, occipital and perisylvian BT features. Sex/handedness differences in BT were found and were related to cognitive sex/handedness differences in verbal-numerical reasoning. We observed differential heritability of up to 56% for BT, especially in temporal language areas, and identified numerous genome- and phenome-wide significant associations pointing to neurodevelopment, cognitive functions, lifestyle, neurological and psychiatric disorders, sociodemographic, cardiovascular and anthropometric traits. This study provides a comprehensive description of BT and insights into biological and other factors that may contribute to the development and individual variations of BT.

## Introduction

The most prominent and earliest reported structural brain asymmetry is an overall hemispheric twist of the brain, known as the Yakovlevian torque^1^. The brain torque (BT) consists of three major components^2^: 1) antero-posterior petalia referring to the protrusion of the right frontal and left occipital poles over the contralateral extremities, 2) left-right tissue distribution asymmetry (TDA) of wider/larger right frontal lobe and left occipital lobe, and 3) bending of the left occipital lobe over the right across the midline skewing the interhemispheric fissure towards the right. With the advance of high-resolution neuroimaging and brain mapping techniques, more complex, localized brain asymmetries have been widely revealed^3, 4^.

Structural brain asymmetries in functionally related brain regions have been associated with functional lateralization of language, handedness and multiple cognitive functions^2, 5^. However, the variability of brain asymmetry, in fact, is affected by various biological, genetic, environmental and clinical factors, which makes the interrelation between structural and functional asymmetry complicated^2, 6^. A number of studies have found differential developmental events of structural brain asymmetries during childhood^7, 8^ and throughout the lifespan^9, 10^. Sex differences in brain asymmetry have been observed in distributed brain regions using diverse brain structural measurements^11^. Furthermore, brain asymmetries have been observed prenatally ^12, 13^, indicating an inherently lateralized, genetically mediated program of brain development^14^. However, recent large-scale imaging genetic studies revealed that heritability of structural brain asymmetries was less than 30% in adult cohorts^4, 15-17^, suggesting existence of environmental modifiers^12^. Moreover, aberrant structural asymmetries have been related to diverse brain disorders, such as schizophrenia^18^, attention-deficit/hyperactivity disorder (ADHD)^7, 19^, autism spectrum disorder (ASD)^20^, dyslexia^21^, Alzheimer’s disease (AD)^22^ and mood disorders^23, 24^. By far, the literature has not been consistent^4, 6, 25^. Most previous studies were based on limited sample sizes and heterogeneous methods for brain asymmetry measurement and analysis. Therefore, it is important to characterize structural asymmetries using large samples and include all relevant structural determinants and possible influencing factors to acquire definitive and normative references for future studies^4, 6^.

The current study was aimed to characterize BT and assess its relationships with potential genetic and nongenetic modifiers in a large sample comprised up to 24,112 individuals from 6 neuroimaging cohorts (Supplementary Table 1): the Adolescent Brain Cognitive Development (ABCD^26^, N=2,769), the Human Connectome Project (HCP^27^, N=860), the International Consortium for Brain Mapping (ICBM^28^, N=371), the Pediatric Imaging, Neurocognition, and Genetics (PING^29^, N=677), the Philadelphia Neurodevelopmental Cohort (PNC^30^, N=922) and the UK Biobank (UKB^31^, N=18,513). For a complete characterization, we systematically quantified all relevant components of BT (Extended Data Fig. 1). Lobar measures of fontal/occipital petalia (protrusion along the antero-posterior axis) and shift (protrusion along the dorso-ventral axis) were computed as the relative displacements of the left and right frontal/occipital extremes. Lobar measures of frontal/occipital bending were computed as the angles between the planes best fitting the interhemispheric fissure in the frontal/occipital regions and the mid-sagittal plane (MSP). Sectional tissue distribution asymmetries (TDA) were measured as left-right differences in hemispheric width (Asym_width_) and perimeter (Asym_perimeter_) in 60 contiguous coronal brain slices. Vertex-wise interhemispheric surface positional asymmetries (SPA) were calculated along the left-right (Asym_LR_), antero-posterior (Asym_AP_) and dorso-ventral (Asym_DV_) axes. We assessed age-related differences in all these profiles across the lifespan (3–81 years), as well as the effects of sex, handedness, sex-by-handedness and total Intracranial volume (TIV). Pedigree- and single-nucleotide polymorphism (SNP)-based heritability of BT features was estimated using available twin and genomic datasets respectively. Genome-wide association study (GWAS) analyses were performed to identify genetic associations with BT at SNP, gene, gene-set and gene-property levels. GWAS summary statistics were also applied to assess genetic correlations of BT with other brain asymmetry related traits. Last, we performed phenome scan (PHESANT^32^) analyses to explore other potential correlates of BT by searching over the extensive phenotypic data from the UKB database, including health and lifestyle questionnaires, sociodemographic factors, physical and cognitive measures, medical imaging markers and etc.

## Results

### Population-level average brain torque

Patterns of BT were demonstrated in the individual and pooled datasets. In accord with the widely reported anticlockwise, antero-posterior torque, the pooled sample showed right-frontal (T=-62.60, P<2.23e-308) and left-occipital (T=-96.48, P<2.23e-308) petalia, leftward-frontal (T=22.98, P=7.33e-116) and rightward-occipital (T=98.32, P<2.23e-308) bending, rightward-frontal and leftward-occipital Asym_width_ and Asym_perimeter_ (P<4.17e-4) (Fig. 1-A and B). In addition, on average, both the frontal (T=-8.51, P<9.40e-18) and occipital (T=-44.29, P<2.23e-308) poles of the left hemisphere were shifted inferiorly relative to the right along the dorsal-ventral axis (Fig. 1-A). We also found that the petalia, bending and shift in the occipital lobe were significantly larger than their frontal counterparts (Supplementary Table 2). The prevalence of these BT patterns demonstrated their predominance in the population (Supplementary Table 3 and Supplementary Fig. 1).

Results of the vertex-wise SPA were well in line with the lobar and sectional data. In the pooled sample, most of the left hemispheric surface was displaced posteriorly, inferiorly and nearer to MSP relative to the right (Fig. 1-C). Additionally, we found that a leftward Asym_LR_ in the superior temporal sulcus (STS) was surrounded by opposite effects and the most pronounced posterior and ventral relative displacements were located in the posterior boundary of the Sylvian fissure (SF).

**Figure 1.**
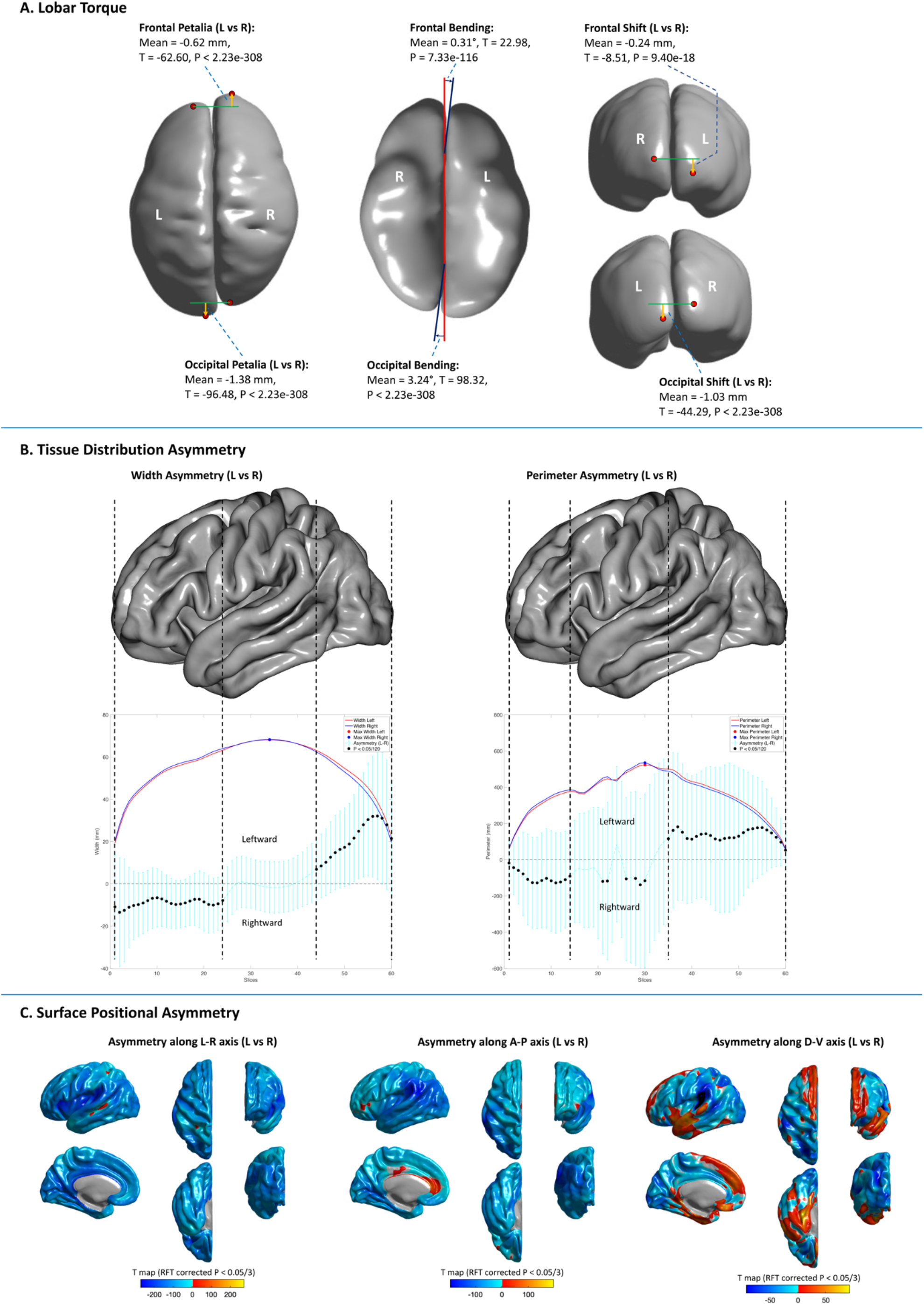
Population-level, average brain torque profiles computed in the pooled sample. (A) Statistics of petalia (left panel), bending (middle panel) and shift (right panel) are illustrated on a smoothed cortical surface model. Red points represent frontal/occipital extremes. Orange arrows show directions of petalia and shift. In the middle panel, red and blue lines respectively represent the middle-sagittal plane (MSP) and the fitted frontal/occipital interhemispheric planes in the axial view. (B) Sectional asymmetries in hemispheric width (left panel) and perimeter (right panel). Significant asymmetries (P < 0.05/120 and asymmetry/(left+right) > 2%) are marked with black dots. In order to show the hemispheric width/perimeter and the asymmetries in the same plot, asymmetry metrics were multiplied by 10. The left hemisphere of the FreeSurfer fsaverage cortical surface template is displayed above the asymmetry plots as the reference of brain anatomy. (C). Significant (random field theory (RFT) corrected P < 0.05/3) surface positional asymmetries along the Left-Right (left panel), Antero-Posterior (middle panel) and Dorso-Ventral (right panel) axes. Color bars represent T statistics. In the left panel, red-yellow and blue-cyan respectively show leftward and rightward displacements of the left hemisphere relative to the right; in the middle panel, red-yellow and blue-cyan respectively show forward and backward relative displacements; in the right panel, red-yellow and blue-cyan respectively show upward and downward relative. L = Left, R = Right, A-P = Antero-Posterior, D-V = Dorso-Ventral.

The population-level, average patterns of BT were largely reproduced in the individual datasets, especially in the ABCD, PING and PNC cohorts composed of children and adolescents and the UKB cohort composed of middle-aged and older adults. Exceptions were observed in the HCP and ICBM cohorts mainly consisting of young adults. In these cohorts, the frontal effects of petalia, shift, bending, Asym_width_, Asym_perimeter_ and/or Asym_AP_ were absent indicating possible age differences (Supplementary Table 2 and Supplementary Fig. 2 and 3).

### Age-related differences in brain torque

Complex (linear and nonlinear) cross-sectional age trajectories of BT profiles were characterized across the lifespan (3-81 years) in the pooled sample. Significant nonlinear (cubic or quadratic) age effects were detected for nearly all the lobar torque measures (Fig. 2-A), except the occipital shift that showed a linear age effect. For sectional TDA, Asym_width_ showed nonlinear age effects in most brain sections (Fig. 2-B). Nonlinear age effects on Asym_perimeter_ were mainly observed in the sections covering the prefrontal cortex (PFC) (Fig. 2-B). For SPA, nonlinear age effects were distributed in most cortical areas for Asym_LR_ and Asym_AP_ and primarily in most of the perisylvian cortex and surrounding areas for Asym_DV_ (Fig. 2-C). It is notable that most of the observed nonlinear age trajectories were characterized by an initial negative age-BT association from childhood to early young adulthood, followed by a positive association in late young adulthood, and then a stabilization of the torque (cubic model) or lack of the phase of stabilization (quadratic model) in late adulthood; most of the linear age trajectories were delineated by a positive BT-age association across the lifespan.

**Figure 2.**
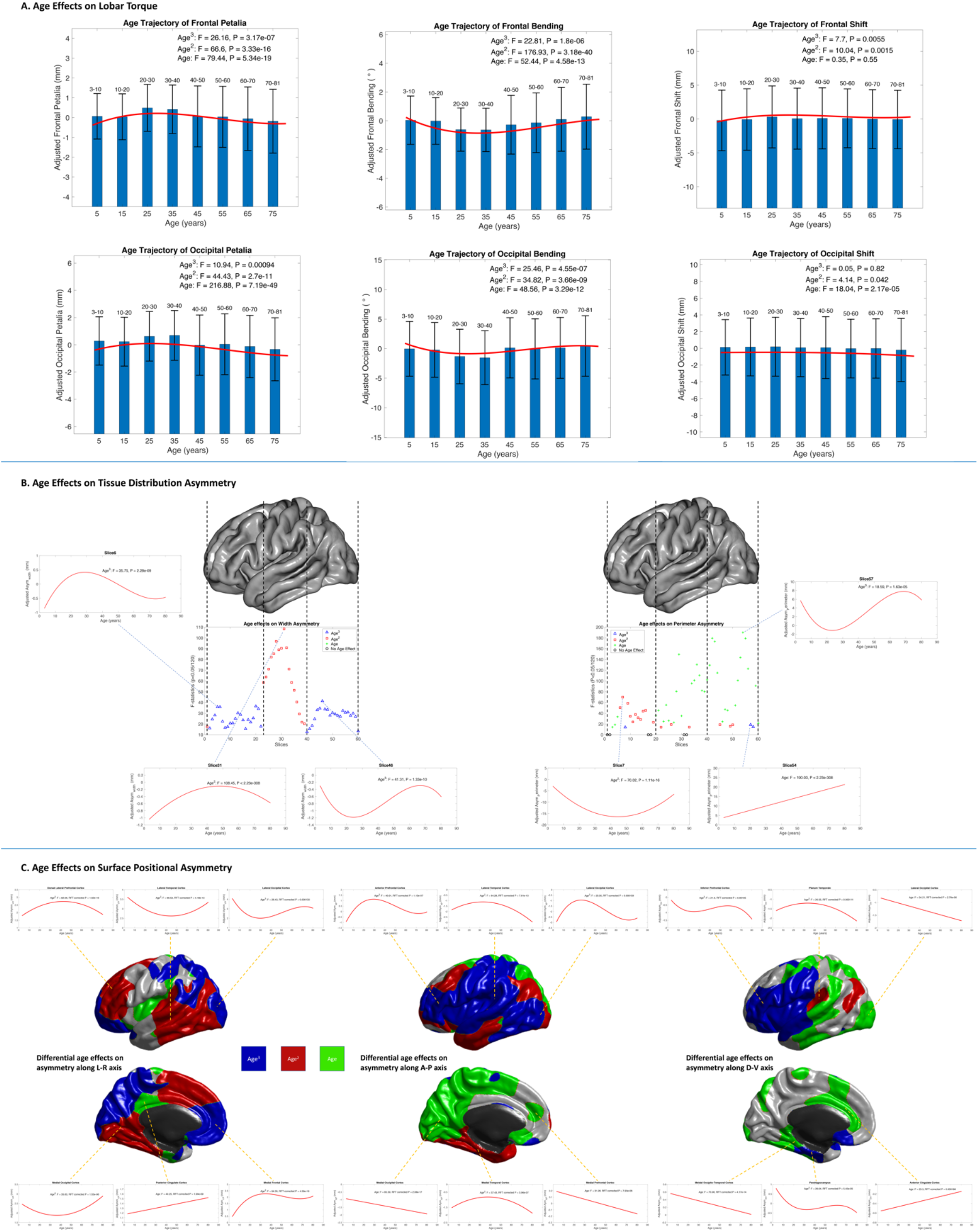
Age-related differences in brain torque profiles. (A) Red lines represent fitted age trajectories of frontal/occipital petalia (left column), bending (middle column) and shift (right column). Statistics for cubic, quadratic and linear age effects are annotated in each plot. Bar graphs with error bars show the means and standard deviations of adjusted BT measures in sub-age groups. (B) Age effects on asymmetries in hemispheric width (left column) and perimeter (right column). Significant (P < 0.05/120) cubic, quadratic and linear age effects are marked with blue triangles, red square and green ‘+’ symbols respectively. Fitted age trajectories are depicted for selected brain slices. The left hemisphere of the FreeSurfer fsaverage cortical surface template is displayed above the plots of age effects as the reference of brain anatomy. Asym_width_ = width asymmetry, Asym_perimeter_ = perimeter asymmetry. (C) Distributions of significant (random field theory (RFT) corrected P < 0.05/3) cubic (blue areas), quadratic (red areas) and linear (green areas) age effects on surface positional asymmetries along the Left-Right (left column), Antero-Posterior (middle column) and Dorso-Ventral (right column) axes. Fitted age trajectories are depicted for selected vertices. L-R = Left-Right, A-P = Antero-Posterior, D-V = Dorso-Ventral. Asym_LR_ = asymmetry along left-right axis, Asym_AP_ = asymmetry along Antero-Posterior axis, Asym_DV_ = asymmetry along Dorso-Ventral axis.

### Sex and handedness effects on brain torque

Compared to females, males showed higher prevalence of the typical right-frontal and left-occipital petalia (males: 58.53% vs females: 54.36%, χ^2^=41.28, P=1.32e-10) and the typical leftward-frontal and rightward-occipital bending (males: 44.66%% vs females: 41.32%, χ^2^=26.55, P=2.57e-7) (Supplementary Table 4 and 5). Consistently, the population-level, average BT patterns were generally enlarged in males relative to females, including larger right-frontal (T=8.27, P=7.06e-17) and left-occipital (T=6.51, P=7.81e-11) petalia (Fig. 3-A), increased leftward-frontal bending (T=6.81, P=4.90e-12) (Fig. 3-A), greater rightward-frontal and leftward-occipital Asym_width_(P<4.17e-4) and leftward-occipital Asym_perimeter_(P<4.17e-4) (Fig. 3-B), while the rightward TDA in the slices corresponding to the planum temporale (PT) and surrounding areas were reduced in males (P<4.17e-4) (Fig. 3-B). Most of the population-level average patterns of SPA were exaggerated in males compared to females as well (Fig. 3-C).

**Figure 3.**
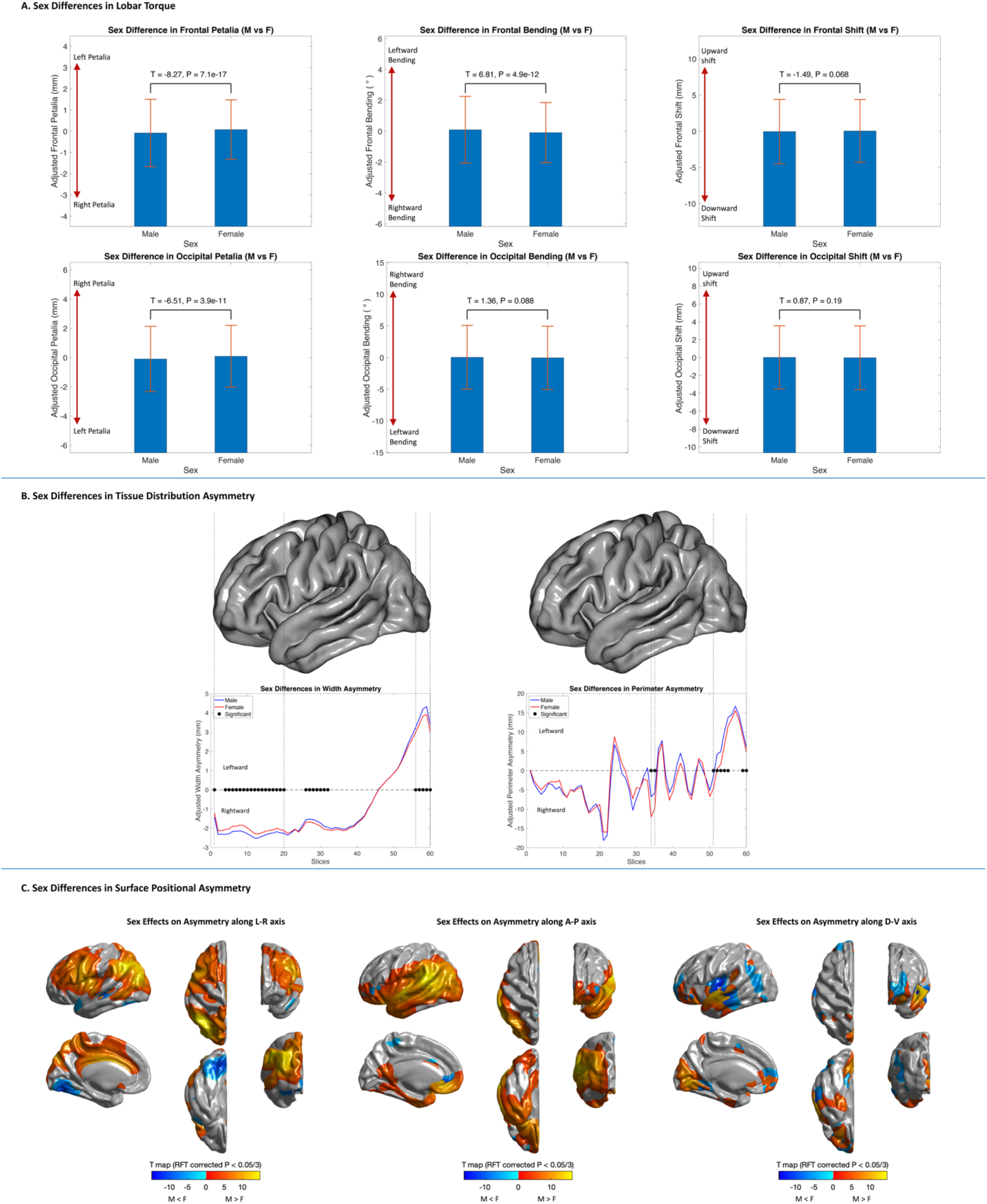
Sex differences in brain torque profiles. (A) Bar graphs with error bars represent adjusted frontal/occipital petalia (left column), bending (middle column) and shift (right column) for males and females. Statistics for handedness comparisons are annotated in each plot. (B) Blue and red lines represent adjusted sectional hemispheric width (left column) and perimeter asymmetries (right column) for males and females, respectively. Black dots indicate significant (P < 0.05/120) sex differences. The left hemisphere of the FreeSurfer fsaverage cortical surface template is displayed above the plots of sex differences as the reference of brain anatomy. (C) Significant (random field theory (RFT) corrected P < 0.05/3) sex effects on surface positional asymmetries along the Left-Right (left panel), Antero-Posterior (middle panel) and Dorso-Ventral (right panel) axes. Color bars represent T statistics. Red-yellow and blue-cyan indicate enlargement and reduction of asymmetry magnitude in males relative to females, respectively. L-R = Left-Right, A-P = Antero-Posterior, D-V = Dorso-Ventral.

Comparing different handedness groups, mixed-handers showed a lower prevalence of the typical right-frontal and left-occipital petalia than right-handers (mixed-handers: 51.09% vs right-handers: 56.74%, χ^2^=10.88, P=9.71e-4), and a higher proportion of right-frontal and right-occipital petalia than left-handers (mixed-handers: 12.03% vs left-handers: 8.56%, χ^2^=8.64, P=0.0033) (Supplementary Table 4 and 5). For BT magnitudes, right-handers showed increased leftward-frontal bending than left-handers (T=4.32, P=7.79e-6) and larger left-occipital petalia than mixed-handers (T=-2.81, P=0.0025) (Fig. 4-A). Right-handers also showed more leftward-occipital Asym_width_than left-handers, and reduced rightward Asym_width_ at the slices covering PTO than left- and mixed-handers (Fig. 4-B). Handedness differences in SPA were also detected between right- and left-handers, primarily in the medial parietal cortex for Asym_LR_, in the insula and perisylvian cortex for Asym_AP_, and in the motor, temporal and lateral occipital cortices for Asym_DV_ (Fig. 4-C). No effects of sex-by-handedness interaction and TIV were found for all BT features.

**Figure 4.**
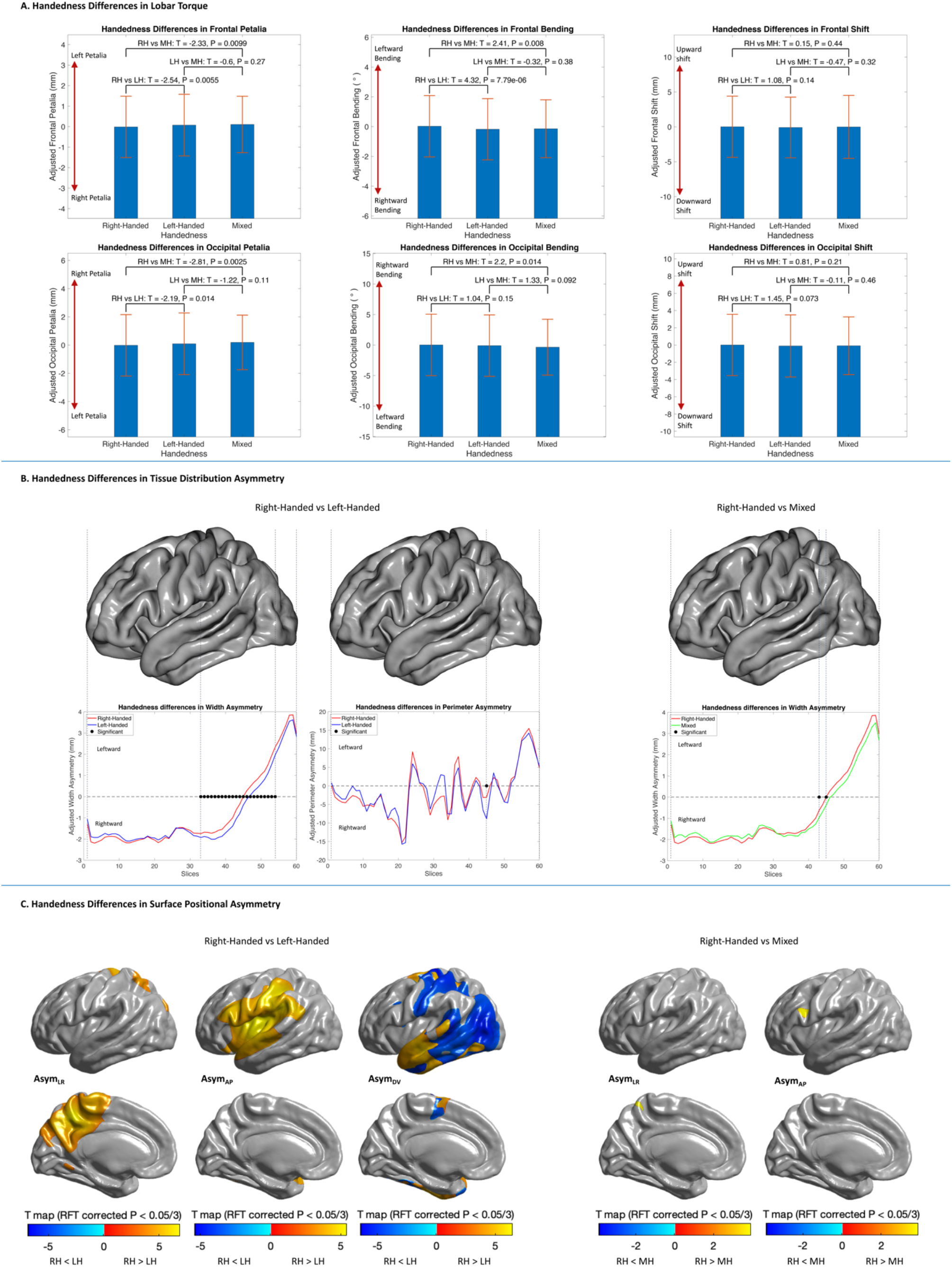
Handedness differences in brain torque (BT) profiles. (A) Bar graphs with error bars represent adjusted frontal/occipital petalia (left column), bending (middle column) and shift (right column) for right-, left- and mixed-handers. Statistics for handedness comparisons are annotated in each plot. (B) Left panel shows differences between right- and left-handers in sectional width and perimeter asymmetries; right panel shows differences between right- and mixed-handers in sectional width asymmetries. Significant (P < 0.05/120) handedness differences were highlighted with black dots. The left hemisphere of the FreeSurfer fsaverage cortical surface template is displayed as the reference of brain anatomy. No significant differences between right- and mixed-handers in perimeter asymmetries and between left- and mixed-handers in width and perimeter asymmetries were found. Thus, the results are not shown here. (C) The left panel shows significant (random field theory (RFT) corrected P < 0.05/3) differences between right-handers (RH) and left-handers (LH) in surface positional asymmetries along the Left-Right (Asym_LR_), Antero-Posterior (Asym_AP_) and Dorso-Ventral (Asym_DV_) axes; the right panel shows significant differences between RH and mixed-handers (MH) in Asym_LR_ and Asym_AP_. Color bars represent T statistics and directions of between-group differences. No significant differences between right- and mixed-handers in Asym_DV_ and between LH and MH in all surface positional asymmetry measures were found. Thus, the results are not shown here.

### Heritability of brain torque

Pedigree-based heritability 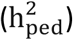 of BT features was estimated using the twin data of the ABCD and HCP cohorts. The ABCD sample showed a 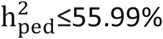 among all BT profiles (Fig. 5),especially for frontal petalia 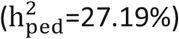, occipital bending 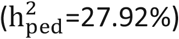, TDA in the posterior portion of the brain 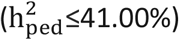, Asym_AP_ in the frontal and occipital regions 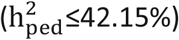, and Asym_LR_ in PT 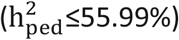. The HCP cohort showed lower heritability estimates than ABCD (Fig. 5). The most heritable profiles were occipital bending 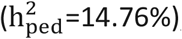, TDA in the anterior and posterior portions of the brain 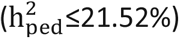, Asym_AP_ in the frontal and occipital regions 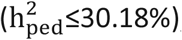, Asym_DV_ in the Heschl’s gyrus 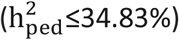, and Asym_LR_ in STS 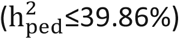. In addition, SNP-based heritability 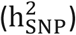 was estimated using the UKB dataset (ABCD, PING and PNC datasets were not used due to insufficient sample sizes, see Methods) and the meta-GWAS summary statistics. We observed patterns that were similar to the pedigree-based results for HCP, however, the SNP-based estimates were remarkably lower than the pedigree-based ones possibly because the GWAS genotyping arrays do not contain all variants in the genome (Supplementary Table 6 and Supplementary Fig. 4 and 5).

**Figure 5.**
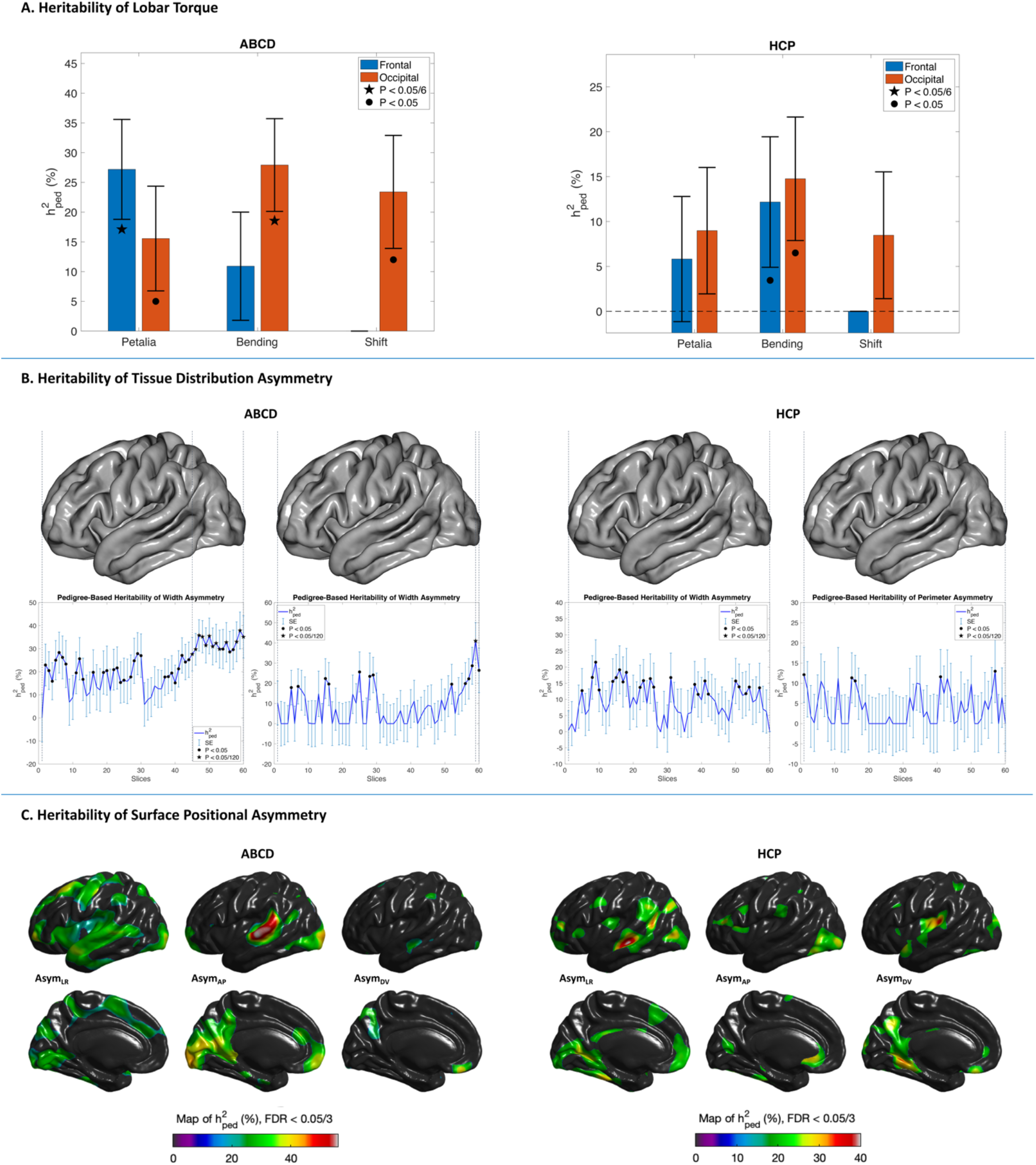
Pedigree-based heritability 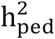 of brain torque profiles estimated using the ABCD (left column) and HCP (right column) datasets. (A) Bar graphs with error bars illustrate 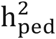estimates and standard errors (SE) for frontal/occipital petalia, bending and shift. Black pentagrams indicate significant heritability (P < 0.05/6). (B) Blue lines with error bars represent 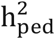estimates and SE for sectional hemispheric width and perimeter asymmetries. Black pentagrams indicate significant heritability (P < 0.05/120). The left hemisphere of the FreeSurfer fsaverage cortical surface template is displayed above the heritability plots as the reference of brain anatomy. (C) Maps of 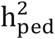estimates (thresholded at FDR < 0.05/3) of surface positional asymmetries along the Left-Right, Antero-Posterior and Dorso-Ventral axes. Asym_LR_ = asymmetry along left-right axis, Asym_AP_ = asymmetry along Antero-Posterior axis, Asym_DV_ = asymmetry along Dorso-Ventral axis.

### Genome-wide association analyses

GWAS analyses were first performed for 6 lobar measures, 60*×*2 sectional measures, and 74*×*3 regional mean SPA measures in the ABCD, PING, PNC and UKB samples individually, and then combined by meta-analysis using METAL with genomic control^33^. We observed no evidence for significant residual stratification effects (for all BT measures, λ_GC_=0.9809 to 1.0195, LDSC Intercept = 0.9824 to 1.0089, LDSC Intercept SE = 0.0038 to 0.0052). At the genome-wide significance threshold of P<5e-8, we identified 133 gene-BT associations, including 1 independent lead SNP associated with frontal bending, 13 independent loci across 9 chromosomes influencing TDA of 14 brain slices, and 72 independent variants across 20 chromosomes associated with the regional mean SPA of 51 brain regions. 2 out of the 133 associations remained significant after correcting for multiple comparisons between BT measures (P<5e-8/348=1.44e-10) (see Fig. 6, Supplementary Table 8 and Supplementary Fig. 6 and 7). Association lookups via the NHGRI-EBI GWAS catalog^34^ showed that 84 out the 86 lead variants were intronic/close to genes that were previously associated with other traits, such as gray matter (GM)/white matter (WM)/ventricular structures, cognitive abilities, neurological and psychiatric disorders, cardiovascular, autoimmune and inflammatory diseases, educational attainment, alcohol consumption, smoking, sleep, anthropometric characteristics and physical appearance (Supplementary Table 9 and 10).

**Figure 6.**
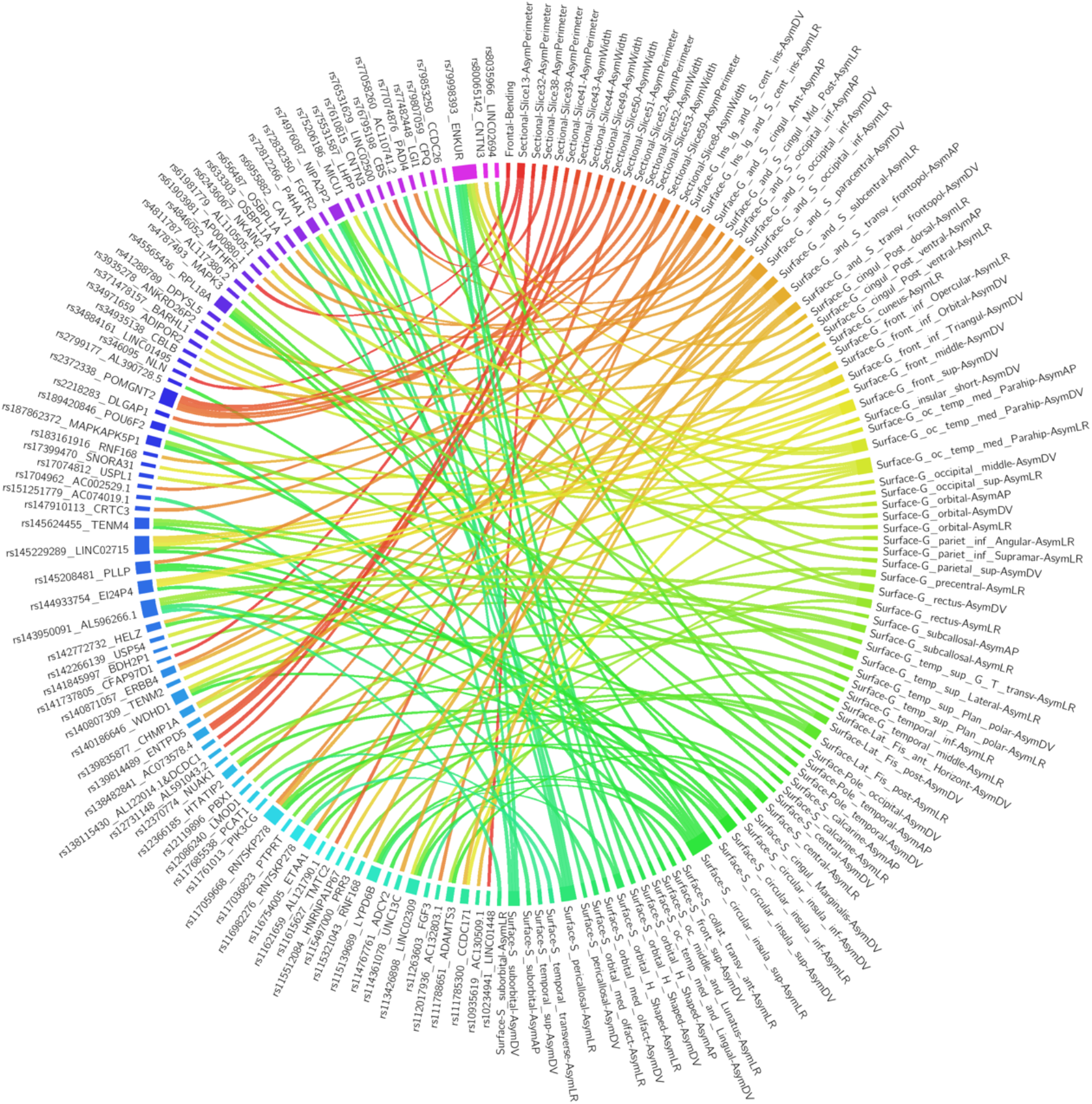
Overview of significant (P < 5E-8) gene-brain torque (BT) associations identified in the meta-GWAS. Ribbons in the connectogram plot illustrate associations of the 86 lead variants (left) with different BT profiles (right). Different colors represent different lead variants or BT profiles. Ribbons are colored with the colors of linked BT profiles. Ribbon thickness represent the -log(P) value of the corresponding association. The lead variants are annotated with SNP IDs and closest genes. WidthAsym = Width Asymmetry, PerimAsym = Perimeter Asymmetry, AsymLR = asymmetry along left-right axis, AsymAP = asymmetry along Antero-Posterior axis, AsymDV = asymmetry along Dorso-Ventral axis. Surf = Surface. For the abbreviations of brain surface regions, see Supplementary Table 7.

### Genome-wide gene-based and gene-level analyses

MAGMA^35^ genome-wide, gene-based analysis on the meta-GWAS results revealed 22 genes associated with occipital bending, TDA of 5 brain slices and regional mean SPA of 38 brain regions, at the significance threshold of P<0.05/18359 genes=2.72e-06 (Supplementary Table 11). Association lookups for these genes were summarized in Supplementary Table 12 and 13. None of the gene-BT associations remained significant when further correcting for the number of BT measures (P>0.05/18359/348=7.83e-9).

MAGMA gene-set analysis showed 75 biological pathways for occipital petalia, TDA of 23 brain slices and mean SPA of 38 brain regions, at the critical threshold of P<0.05/7563 gene sets=6.61e-6 (Supplementary Table 14). Among them were the Gene Ontology (GO) biological process (BP) terms ‘BP:GO_neuronal_action_potential_propagation’ associated with mean Asym_AP_ of the fusiform gyrus, and ‘BP:GO_regulation_of_neuron_projection_development’, ‘BP:GO_neuron_differentiation’, ‘BP:GO_neuron_development’ and ‘BP:GO_cell_morphogenesis_involved_in_neuron_differentiation’ associated with the mean Asym_DV_ of STS, and ‘BP:GO_central_nervous_system_myelin_maintenance’ associated with the mean Asym_DV_ of the lateral orbital sulcus. None of the 75 enrichments would be significant with further correction for the number of BT measures (P>0.05/7563/348=1.90e-9).

MAGMA gene-property analysis testing the genome-wide, gene-based analysis results with respect to the BrainSpan^36^ age-specific gene expression data (see Supplementary Table 15 and Supplementary Fig. 8 and 9) showed that gene expression levels during the first trimester and early second gestational trimester (8–16 postconceptional weeks (pcw)) were positively correlated with the genetic associations with Asym_width_ of brain slices covering PT, PTO and the anterior PFC and with mean Asym_AP_ and/or Asym_LR_ in multiple regions of temporal, occipital, circular insular and precentral cortices (P<0.05/31 age stages=0.0016). We also observed that higher gene expression levels were predictive of the genetic associations with frontal bending and frontal Asym_perimeter_ throughout most of the neurodevelopmental stages from middle- or late-prenatal stages (24 and 37 pcw) until late young adulthood (30-40 years) (P<0.05/31=0.0016). These gene-property results did not remain significant after further correcting for the number of brain torque measures (P>0.05/31/348=4.63e-6).

### Genetic correlation with other traits

We performed genetic correlation analyses with our meta-GWAS results in relation to other complex traits that have been associated with structural and/or functional brain asymmetries (see Methods). No genetic correlation was found at the Bonferroni-corrected critical threshold. We observed trends (P<0.05) of genetic correlations with ADHD and/or AD for TDA of multiple slices corresponding to PT and surrounding areas, and with educational attainment, intelligence, AD, schizophrenia and/or bipolar disorder for mean Asym_LR_ and/or Asym_DV_ in several brain regions of the orbitofrontal, insular, parietal, temporal and occipital cortices (Supplementary Figure 10 and Supplementary Table 16).

### Phenome scan analyses

We systematically tested correlations of BT features with the extensive phenotypes from UKB using the Phenome Scan Analysis Tool (PHESANT^32^). 942 BT-related phenotypes were identified at the significance level adjusting for both the number of phenotypes and of BT measures (P<0.05/6334/348=2.27e-8). The majority (832 out of 942) were regional brain MRI measures, including regional volume, surface area and mean thickness, microstructural measures of WM tracts, and volume, mean intensity and/or T2^*^ of subcortical regions and lateral ventricles (Fig. 7). Notably, the associations with these cortical morphometries and WM microstructures displayed interhemispheric asymmetries and anterior-posterior twist, i.e., the association with a lateral regional MRI variable was always reversed or absent in its counterpart in another hemisphere, and the directions of associations with lateral frontal regions were generally homogeneous with occipital regions in another hemisphere rather in the same hemisphere (Fig. 8). Moreover, among the identified phenotypes, the most pronounced associations (P<5e-324) were observed for the mental health variable of bipolar and major depression status, the lifestyle variable of smoking status and the cognitive function of verbal-numerical reasoning (fluid intelligence) (Fig. 7). The other associated phenotypes included lifestyle behavior of alcohol consumption, sociodemographic factors of education, ethnicity and household, physical measures (e.g. body mass index (BMI), height and weight and etc.), and other variables e.g., country of birth, home area population density, number of full sisters and natural hair color and etc. Additional 308 phenotypes were detected at the significance level controlling for only the number of phenotypes (P<0.05/6334=7.89e-6), including cognitive function of prospective memory, circulatory system diseases (e.g. ICD10 code: I10 essential hypertension and I73.9 peripheral vascular disease), and other nonbrain health-related outcomes. Detailed results of the PHESANT scans are listed in Supplementary Table 17-22.

**Figure 7.**
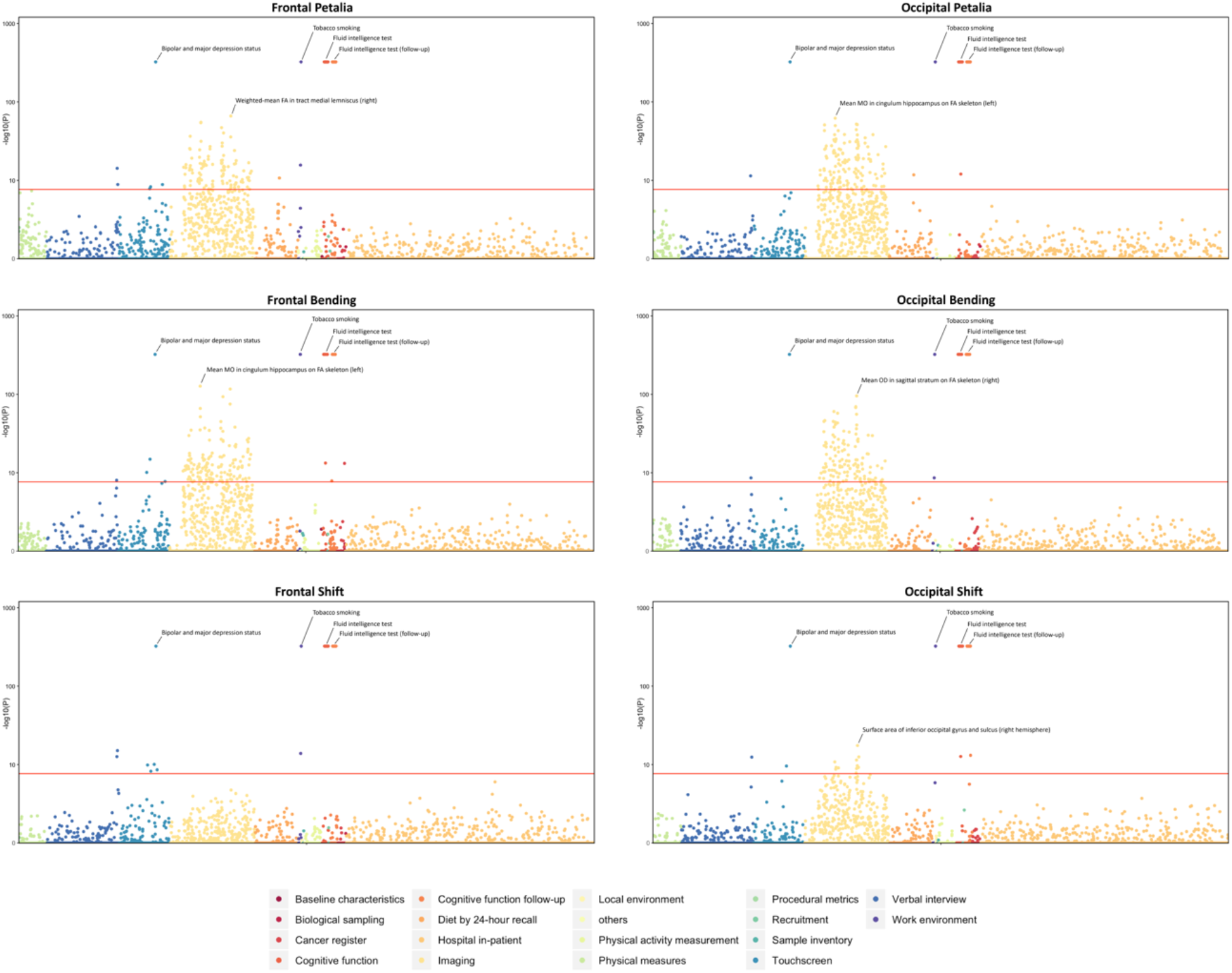
Phenome-wide association scans for lobar brain torque profiles. Manhattan Plots for phenome-wide associations of frontal/occipital petalia, bending and shift. Red lines indicate the phenome-wide significant threshold adjusted for the number of brain torque measures (P < 0.05/6334/348 = 2.27e-8). For detailed results of phenome-wide association scans, see Supplementary Table 17-22.

**Figure 8.**
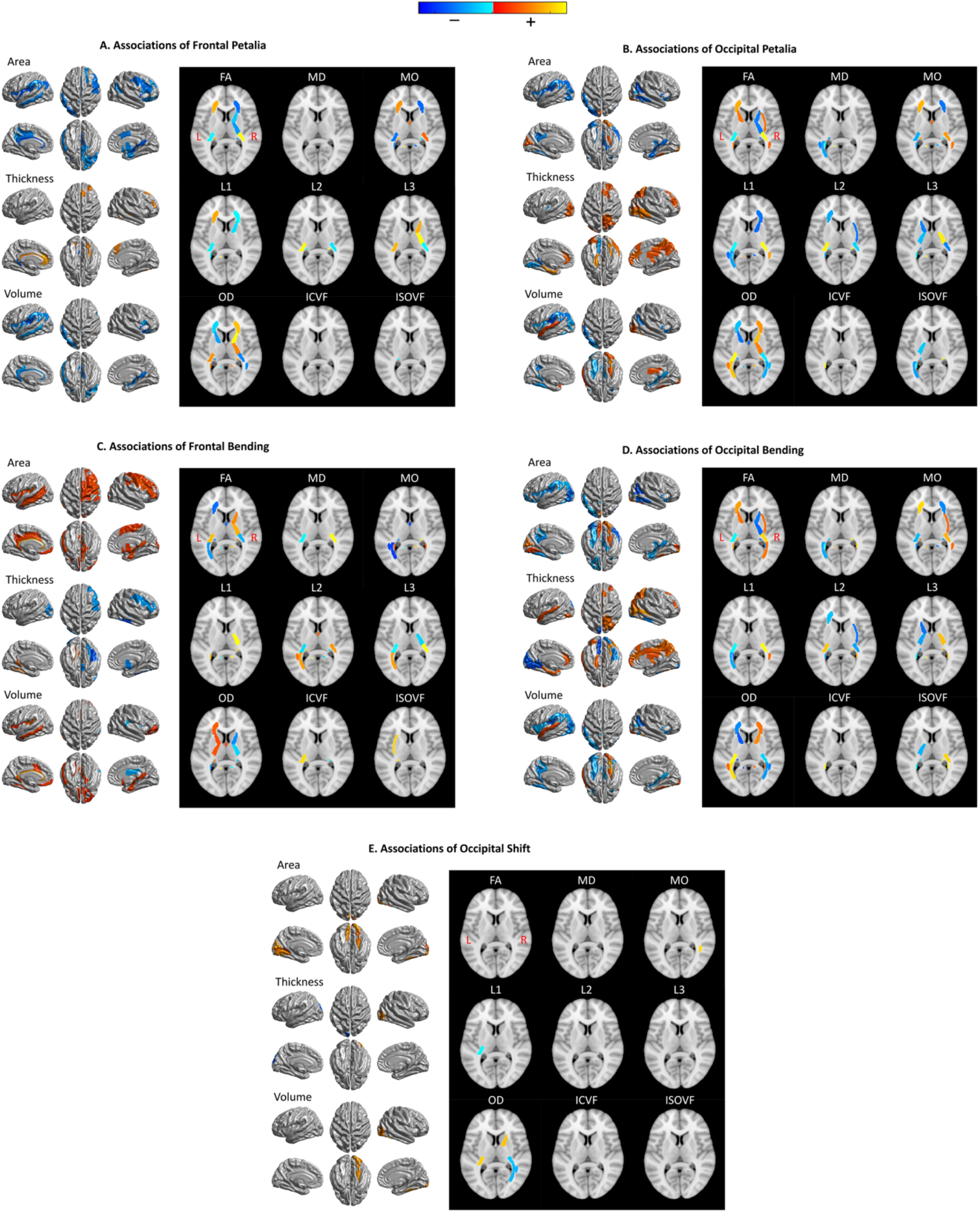
Maps of associations between lobar brain torque (BT) profiles and gray matter (GM) and white matter (WM) metrics. The panels show phenome-wide significant associations (P < 0.05/6334/348 = 2.27e-8) of frontal petalia (A), occipital petalia (B), frontal bending (C), occipital bending (D) and occipital shift (E). No significant associations between frontal shift and brain imaging metrics were found. The results, therefore, are not shown here. Red-yellow and blue-cyan represent positive and negative associations respectively. On the left of each panel, statistics of associations between BT and regional measures of cortical surface area, thickness and volume are rendered on the FreeSurfer fsaverage cortical surface template; on the right, association statistics for WM tracks are displayed on a selected axial slice (z = 8) of the MNI152 brain template. FA = fractional anisotropy, MD = mean diffusivity, MO = mode of anisotropy, L1/ L2/L3 = the three eigenvalues of diffusion, OD = orientation dispersion, ICVF = intra-axonal volume fraction, ISOVF = isotropic volume fraction. For detailed results of phenome-wide association scans, see Supplementary Table 17-22.

## Discussion

We presented a comprehensive, systematic study of human BT components in a large sample comprised up to 24,112 individuals from 6 cohorts. We demonstrated the anticlockwise torque as the population-level, average pattern. Many BT features showed a U-shaped lifespan age trajectory with a breakpoint in the late young adulthood and were associated with sex and handedness. Our heritability analysis revealed differential heritability for multiple BT profiles, as well as a possible age difference in the heritability. The GWAS and PHESANT analyses identified a number of independent genetic variants and complex phenotypic variables that were correlated to BT, which together may indicate an interrelated asymmetric system of the human body.

### Human brain torque

The current study represents the first, large-scale, systematic study of all BT components. We quantified complex BT features in a large sample using a framework integrating a set of automatic 3D brain shape analysis approaches. The lobar and sectional measures provided direct modeling of the primary BT components^2^ as well as frontal bending and dorsal-ventral shift that were rarely studied previously^37^. The vertex-wise analysis yielded detailed, comprehensive measurements of SPA in three orthogonal directions^38^. These methods have been reliable and robust in several recent, small sample-based studies on species differences in BT between human and chimpanzees^37-40^. The population-level average BT patterns were largely consistent between the cohorts employed here and also were homogeneous with those reported in the earlier studies^37-40^, which further demonstrates the reliability and robustness of these algorithms.

The present large-scale data confirmed the predominance of the anticlockwise antero-posterior twist patterns of the brain in the population, comprising right-frontal and left-occipital petalia, leftward-frontal and rightward-occipital bending, and rightward-frontal and leftward-occipital TDA, as well as an anticlockwise twist along the dorsal-ventral axis. These typical torque patterns were well reproduced in vertex-wise analyses of SPA. In addition, the SPA data showed a leftward Asym_LR_ (left surface is farther away from MSP than right) in STS surrounded by opposite effects, indicating a greater STS depth in the right hemisphere than in the left. Relative to the right, the posterior boundary of the left SF displayed pronounced posterior and ventral displacements and the left lateral superior temporal gyrus was lifted upward, implying that the left SF is longer, narrower and less curved than the right. It has recently been illustrated that the typical BT patterns together with the STS and SF asymmetries were absent in chimpanzees^37-41^, suggesting an important role of BT in human brain evolution.

Compared to females, the typical BT patterns were more predominant in males in terms of population prevalence and magnitude. This finding is consistent with the accumulative evidence suggesting that females have a more bilateral brain organization than males^11^. Right-handedness was associated with multiple distinct BT patterns compared to mixed- and left-handedness. The relationship between sex and handedness differences in brain asymmetry and cognitive abilities has not been revealed yet^11, 42^. Our PHESANT analysis identified the cognitive function of verbal-numerical reasoning as one of the most prominent phenotypic variables associated with BT. We also found a significantly increased verbal-numerical reasoning score in males compared to females (cohen’s d=0.14, P=3.94e-23) and in right-handers compared to mixed-handers (Cohen’s d=0.14, P=0.01) (Supplementary Fig. 11). Taken together, the current neuroimaging and cognitive data may reflect potential relations between the sex dimorphisms and handedness differences (right-handed vs mixed-handed) in BT and in the ability of verbal-numerical reasoning.

Recently, Kong and colleagues^16^ performed the first large-scale study of global human BT using overall brain skew parameters derived from MRI registration. The current results of complex BT components are largely consistent with their brain skew data, and provide details about how the reported anticlockwise antero-posterior and dorsal-ventral skew are formed.

### Development of brain torque

To date, the developmental mechanism of brain asymmetry has been unclear. In line with the brain skew study^16^, here, BT was systematically associated with diverse brain MRI measures in widespread cortical/subcortical regions, WM tracts and lateral ventricles. In particular, these associations were always lateralized to one hemisphere or reversed between the two hemispheres and were twisted along the anterior-posterior axis. Thus, we speculate that BT development involves lateralized dynamics of distributed GM structural changes^7, 8^, synaptic pruning^24^, axon tension^43^, and/or ventricular enlargement^23^.

The present study illustrated U-shaped, cross-sectional age trajectories peaking in the late young adulthood for many BT features, such as the frontal/occipital petalia, bending and TDA, the frontal shift and the SPA in the frontal, occipital and perisylvian cortices, suggesting a general “first attenuating, then enlarging” dynamic with age. A longstanding 3D lateralized neuro-embryologic growth model proposed by Best^44^ predicted that the early-developed, rightward morphological asymmetry in frontal-motor regions may become attenuated by the later left-biased growth of prefrontal tertiary association regions, whereas the leftward posterior asymmetry may converge with the left-biased growth of posterior tertiary association areas, resulting a more striking left-occipital petalia than right-frontal petalia in adults. In line with this prediction, we observed larger occipital torque than frontal torque. However, the “first attenuating, then enlarging” age trajectory was observed in both frontal and occipital regions, though relatively less pronounced in the posterior region. Recent brain morphometric data has revealed that the development of tertiary association areas is not always left-biased, e.g. cortical thinning is right-biased in PTO^9, 10^. Thus, we adjust and extend Best’s prediction as that during neurodevelopment, the early gross asymmetries may be attenuated by the later growth of tertiary association areas that is biased to the opposite side and the attenuation is more prominent in the anterior regions; then, according to the “first in, last out” theory^45^, the attenuated gross asymmetries in adults may be enlarged by the lateralized, earlier aging of certain tertiary association areas^46, 47^.

We also observed significant heritability for many BT features. The highest heritability was detected in the temporal language areas, including Asym_LR_ in PT for ABCD 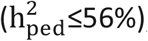; Asym_DV_ in the Heschl’s gyrus 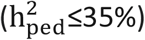 and Asym_LR_ in STS 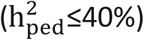 for HCP; and Asym_width_ insections corresponding to PT for UKB 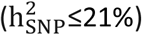. The heritability estimated from the HCP and UKB adult cohorts were in line with the previously reported heritability of structural asymmetries in the same or other adult cohorts^4, 15-17^. Of note, these estimates for adults were generally lower than those computed using the ABCD children sample who likely underwent less environmental influence than adults. This age difference in heritability estimates supports the hypothesis of an environmental contribution to the development of structural brain asymmetry^2, 5^. Consistently, our PHESANT analysis associated BT with various environmental factors, such as lifestyle (tobacco smoking and alcohol consumption), sociodemographics (e.g. education and household), country of birth, and home area population density. Many of these phenotypes were also associated with brain skew using the UKB dataset in ^16^.

### Genetic architecture of brain torque

The meta-GWAS identified 86 independent lead SNPs (LD r^2^<0.2) at the genome-wide significance threshold of P<5e-8. The genome-wide, gene-based analysis revealed additional 22 genes (P<0.05/18359 genes=2.72e-06). It is out of our scope to discuss each of the associations. However, broad patterns emerged showing that the BT-associated genes were also often implicated in the development of cortical/subcortical and ventricular morphology and WM microstructures (e.g. NUAK1, LYPD6B, LINC02694) as well as cognitive abilities and intelligence (e.g. ADCY2, AL596266.1, TENM4). These genes determine development of not only the brain but also susceptibility or resistance to diverse insults: neurodevelopmental (e.g. LINC02715, TENM4), neurodegenerative (e.g. NKAIN2, TENM4), mental (e.g. LINC02694, TENM4), autoimmune (e.g. LINC02694, PCAT1), inflammatory (e.g. CRTC3, CCDC26) and cardiovascular (e.g. MTHFR, AL122014.1), and have been associated with sociodemographic factors (e.g. PTPRT and TENM4 for educational attainment), lifestyle behaviors of alcohol consumption, smoking, sleep and related disorders (e.g. TENM2, AC132803.1), physical characteristics (e.g. ADAMTS3, CCDC171) and physical appearance measurements of eye and hair colors and facial morphology (e.g. PTPRT, CCDC26). There is a wealth of information in the Supplementary Table 8-13 that can be mined for a better understanding of these associations. Of note, the traits that have been associated with the identified genes were largely homogeneous with those BT-associated phenotypes identified in the PHESANT analysis (Supplementary Table 20-22). The current GWAS data, thus, provides a genetic basis for the phenomic associations of BT. Moreover, it has been known that brain asymmetry applies to cognition, emotion, and regulation of autonomic physiological processes, and neurotransmitter, neuroendocrine, immunomodulatory, cardiovascular and vasomotor activity are all asymmetric^48^. Thus, our genomic and phenomic findings may suggest a complex asymmetric system consisting of interrelated components of (neuro)anatomical, neurochemical, physiological and behavioral asymmetries. Additionally, recent large-scale GWASs have associated cytoskeletal-related genes with the global skew^16^ and regional morphometric asymmetries^17^ of the human brain. It has been known that cytoskeleton plays an important role in embryonic development of organ laterality in other species^49, 50^. Here, we also identified 3 key genes in cytoskeletal organization and its regulation: LMOD1, CHMP1A and MAPK3. This finding, together with the earlier GWAS data^16, 17^, evidences existence of a cytoskeleton-mediated mechanism for human brain asymmetry development.

The gene-set enrichment analysis of the GWAS summary statistics revealed that some genes influencing BT were involved in biological processes related to neuronal development and central nervous system myelin maintenance. This may be related to the mechanisms of synaptic pruning^24^ and axon tension^43^ in the development of BT. The gene-property analysis correlated genetic associations with Asym_width_, Asym_AP_ and Asym_LR_ in multiple brain regions with higher expression levels during the first and early second gestational trimester, reflecting genetic contributions to the early establishment of these BT features. The genetic associations with frontal bending and frontal Asym_perimeter_ were correlated with higher gene expression levels since middle- or late-prenatal stages until late young adulthood. This indicates that these frontal torque features may be initiated comparatively later in gestation than Asym_width_ and SPA, while the related genetic mechanism continuously contributes to the development of them.

## Limitations

There are limitations of the current study that should be addressed in future investigations. First, most current results of genetic associations survived in the correction for multiple tests across the genome only but not in the further adjustment for the number of BT profiles, reflecting an insufficient power of the current GWAS. However, the Bonferroni correction for the number of BT features might be overly conservative since many of them are likely correlated. A method that could address the correlation between tests is more appropriate. Second, the majority of individuals included in this study are of European ancestry (∼90%). Especially, only subjects of European ancestry were included in the GWAS analysis in order to avoid false discoveries due to population stratifications. Such limited race focus may raise the question that to what degree our findings can be generalized and applied on global populations. Potential race differences in BT were indicated by the associations with ethnicity and country of birth detected in the PHESANT analysis, though the race distribution was dramatically skewed. It could be an interesting future work to integrate large European and non-European imaging genomics datasets into a global study so as to identify cross-population and population-specific components of brain asymmetry and their genetic correlates.

## Conclusions

This large-scale study illustrated that the anticlockwise BT is predominant in the population. Many frontal, occipital and perisylvian BT profiles showed a U-shaped cross-sectional age trajectory across the lifespan, indicating a “first attenuating, then enlarging” developmental dynamic. Merging neuroimaging and cognitive data, we found possible interrelationships between BT, sex/handedness and cognitive ability of verbal-numerical reasoning. PHESANT analyses associated BT with various GM, WM and ventricular properties as well as other phenotypic variables of cognitive functions, lifestyle, mental health, sociodemographic, cardiovascular and anthropometric traits. The current GWAS analysis identified a number of BT-associated genomic loci, which were associated with many of those BT-associated phenotypes previously, and implicated cytoskeleton in human BT development. Gene-level analyses showed that the genes influencing BT were involved in biological processes of neurodevelopment and they may contribute to the prenatal establishment of BT. In summary, this study yields insights into the biological mechanisms determining the development and individual variability of BT.

## Online methods

### Study Population

The current study utilized neuroimaging data of up to 24,112 subjects from 6 cohorts: the Adolescent Brain Cognitive Development (ABCD), the Human Connectome Project (HCP), the International Consortium for Brain Mapping (ICBM), the Pediatric Imaging, Neurocognition, and Genetics (PING), the Philadelphia Neurodevelopmental Cohort (PNC) and the UK Biobank (UKB). Supplementary Table 1 provides population characteristics of each cohort. All study procedures were approved by the institutional review boards of the participating institutions of these projects, and all participants provided written informed consent

#### Adolescent Brain Cognitive Development

The Adolescent Brain and Cognitive Development Study (ABCD)^26^is a multi-site, longitudinal neuroimaging study following 9–10 year-old youth through adolescence. The ABCD study team employed a rigorous epidemiologically informed school-based recruitment strategy, designed with consideration of the demographic composition of the 21 ABCD sites. This study employed the baseline structural brain MRI dataset of 2,891 subjects from the release 1.0. Details about MRI acquisition protocols are available in^51^. Parental informed consent and child assent were obtained from all participants and approved by centralized and institutional review boards at each data collection site. We discarded 12 subjects due to failed image processing or incomplete metadata and other 110 subjects who were identified as outliers in terms of BT measures, resulting in a sample of 2,769 subjects (age range 9 – 11 years, 1,449 males).

Handedness was evaluated using the ABCD Youth Edinburgh Handedness Inventory (EHI) Short Form^52^ and the sample were classified into left-handed (N = 196), right-handed (N = 2,181) and mixed-handed (N = 392). Details about the ABCD cohort are available at https://abcdstudy.org.

#### Human Connectome Project

The Human Connectome Project (HCP)^27^ is a project to map the neural pathways that underlie brain function and behavior using high-quality neuroimaging data (https://humanconnectome.org/). The details about the HCP dataset are available in the HCP reference manual (https://www.humanconnectome.org/study/hcp-young-adult/document/900-subjects-data-release). All subjects provided written informed consent on forms approved by the Institutional Review Board of Washington University. This study relied on 897 HCP subjects with structural brain imaging from the HCP S900 release (age range 22-37 years, 393 males). In HCP, the strength of hand preference had been assessed with EHI^52^, resulting in scores ranging from - 100 (strong left-hand preference) to 100 (strong right-hand preference). According to the EHI^53^, subjects were classified into left-, right- and mixed-handers based the scores (left: -100 to -60, N = 46; mixed: -60 to 60, N = 195; right: 60 to 100, N = 656). 37 HCP subjects identified as outliers in terms of BT measures were discarded in the current study.

#### International Consortium for Brain Mapping

The International Consortium for Brain Mapping (ICBM) project developed a probabilistic reference system for the human brain^28^. We included structural MRI scans of 641 subjects from the ICBM database. 229 subjects for whom MR image processing was failed and 41 subjects who were outliers in terms of BT measures were discarded in the current study, leaving 371 subjects (age range 18 – 80 years, 197 males). Handedness was quantified with a Laterality Quotient (LQ) using EHI^52^ and the sample were classified into left-handed (N = 34), right-handed (N = 333) and mixed-handed (N = 4). All subjects gave informed consent according to institutional guidelines. Details about the dataset are available at https://ida.loni.usc.edu/.

#### Pediatric Imaging, Neurocognition, and Genetics

The Pediatric Imaging, Neurocognition, and Genetics (PING) data repository is a resource of standardized and curated data of brain MRI, genomics, and developmental and neuropsychological assessments for a large cohort of developing children aged 3 – 20 years^29^. Participants and their parents gave their written informed consent or assent to participate in study procedures. Written parental informed consent was obtained for all PING subjects below the age of 18, and child assent was also obtained for all participants between the ages of 7 and 17. Written informed consent was obtained directly from all participants aged 18 years or older. Detailed information about the PING dataset is available at https://chd.ucsd.edu/research/ping.html and in ^29^. This study utilized structural brain MRI scans of 693 PING participants. We removed 3 participants due failed MRI processing and other 23 participants who were outliers in terms of BT measures, resulting a sample of 677 subjects (age range 3 – 21 years, 355 males, 70 left-handers, 577 right-handers, 30 mixed-handers).

#### Philadelphia Neurodevelopmental Cohort

The Philadelphia Neurodevelopmental Cohort (PNC) is a large-scale initiative that seeks to describe how genetics impact trajectories of brain development and cognitive functioning in adolescence, and understand how abnormal trajectories of development are associated with psychiatric symptomatology^30, 54^. Signed informed consent or assent and parental consent (for participants under age 18) were obtained for all participants. Of the initial 1,445 subjects who completed imaging, 448 subjects were excluded because of a history of potential abnormal brain development or a history of medical problems that may affect the brain. We further excluded 20 subjects with failed MRI processing and 55 subjects as outliers of BT measures, leaving 922 subjects (age range 8 – 23, 428 males, 126 left-handers, 796 right-handers). Details about the PNC dataset are available at https://www.med.upenn.edu/bbl/philadelphianeurodevelopmentalcohort.html.

#### UK Biobank

UK Biobank (UKB) is a large-scale biomedical database and research resource, containing in-depth genetic and health information from about 500,000 UK participants. The present analyses were conducted under UKB application number 25641. Ethical approval was obtained from the research ethics committee (REC reference 11/NW/0382). All participants provided informed consent to participate. Further information on the consent procedure can be found at http://biobank.ctsu.ox.ac.uk/crystal/field.cgi?id=200. This study used brain MRI imaging data^55, 56^ (UK Biobank data-field: 110) from the 2018 August release of 22,392 participants (http://biobank.ctsu.ox.ac.uk/crystal/label.cgi?id=110). Details of the MRI acquisition is described in the UK Biobank Brain Imaging Documentation (http://biobank.ctsu.ox.ac.uk/crystal/refer.cgi?id=1977) and in a protocol form (http://biobank.ctsu.ox.ac.uk/crystal/refer.cgi?id=2367). This study discarded 1,002 participants whose MRI scans did not pass manual quality assessment, 1,148 participants due to data withdraw from UKB or failed image processing, and other 1719 participants who were identified as outliers in terms of BT measures, resulting a sample of 18,513 subjects with age range from 44 to 81 years, and 8,908 male subjects. Handedness was assessed based on responses to the question (UK Biobank data-field: 1707): “*Are you right or left handed?*” with four response options: “*Right-handed*”, “*Left-handed*”, “*Use both right and left equally*”, and “*Prefer not to answer*”. Those who preferred not to answer were excluded for association analysis with handedness, leaving 16,489 right-handers, 1747 left-handers, and 277 mixed-handers with BT measures. We also employed genome-wide genotyping data as described at http://www.ukbiobank.ac.uk/scientists-3/genetic-data/, as well as other phenotypic data, including brain MRI measures of GM morphologies, WM microstructures, T2*, and task-based BOLD (blood-oxygen-level dependent) effects^55, 56^, questionnaires of health and lifestyle, sociodemographic factors, physical and cognitive measures, and etc. In addition to the above exclusion criteria, 795 Participants who reported a diagnosis of neurological and/or psychiatric disorder at scanning (UKB data-field: 20002) were discarded in the analysis of population-level average torque patterns and the effects of age, sex, handedness and TIV (excluded disorders are listed in Supplementary Table 23). These participants were included in the phenome-wide scans for exploring possible associations with clinical traits.

### MRI processing for BT measurements

Different BT components were measured in the 6 neuroimaging datasets using a framework integrating a set of automatic 3D brain shape analysis approaches introduced in recent studies^37-40^ (Extended Data Fig. 1).

All MR images were first preprocessed using the FSL software^57^ v5.0 (https://fsl.fmrib.ox.ac.uk/fsl/fslwiki/) to align brain volumes to the standard MNI template using 7 degrees of freedom transformations (i.e., 3 translations, 3 rotations, and 1 uniform scaling) (Extended Data Fig. 1-1). This preprocessing normalized individual brains into a common coordinate system without distorting the morphological shapes. Next, the normalized images were processed using the FreeSurfer software package^58^v6.0 (https://surfer.nmr.mgh.harvard.edu) to reconstruct cortical hemispheric surfaces (Extended Data Fig. 1-2). The FreeSurfer workflow includes motion correction and averaging of volumetric T1-weighted images^59^, removal of non-brain tissue^60^, automated Talairach transformation, brain volume segmentation^61, 62^, intensity normalization^63^, tessellation of the boundary between gray matter (GM) and white matter (WM), automated topology correction^64^ and surface deformation following intensity gradients to optimally place the GM/WM and GM/cerebrospinal fluid borders at the location where the greatest shift in intensity defines the transition to the other tissue class^65^. Each hemispheric GM and WM surface is composed of 163,842 vertices arranged as 327,680 triangles. Once the surface models are complete, a number of deformable procedures were performed for further data processing and analysis, including surface inflation, registration to a spherical atlas using individual cortical folding patterns to match cortical geometry across subjects^66^.

Accurate definition of the mid-sagittal plane (MSP) is essential for the accuracy of BT measurements, because computation of the angle of interhemispheric fissure bending and surface positional asymmetry replies on it. As a result of the linear spatial normalization using FSL, the x, y and z axes of the MNI coordinate system by default correspond to the left-right, anterior-posterior and dorsal-ventral directions of the brain and the plane x=0 represents the MSP. However, the linear registration is often insufficient to align the true brain MSP to x=0 due to the asymmetric nature of the brain^18, 67^. To address the potential deviation, the MSP was defined as the least squares plane that best fits the vertices on the two hemispheric medial surfaces (excluding the frontal and occipital poles where the interhemispheric fissure bending is maximum) lying within 5 mm to x=0 in the MNI space. Then, the brain orientation was refined by rotating the brain surface with an 3D angle between the plane x=0 and the estimated MSP (Extended Data Fig. 1-4).

To measure gross lobar torque features, a morphologic closing operation was applied to the pial surfaces of smoothed hemispheric volumes in order to fill the sulci (https://surfer.nmr.mgh.harvard.edu/fswiki/LGI) (Extended Data Fig. 1-5). Petalia and shift were computed as the respective displacements of the left and right frontal and occipital extreme points along the antero-posterior and dorso-ventral axes respectively (Extended Data Fig. 1-6). In^37, 40^, to measure frontal and occipital bending of the interhemispheric fissure, planes were first fitted to the vertices of the medial surfaces of the left and right hemispheres in the first (frontal) and last (occipital) quarters of the anterior-posterior length of the brain, next angles between the fitted planes’ normals and the normal of the MSP were calculated for the left and right hemispheres separately, then the angles were averaged between the two hemispheres as the measure of bending. Considering the uncertainty of the normal directions of the fitted planes, such normal-based measure could not directly reflect the true bending in the left or right direction. Thus, we measured the bending with a different procedure: 1) extract the points with the shortest distances to both the left and right hemispheric surfaces, 2) fit planes to these points in the frontal and occipital regions, 3) the frontal and occipital bending was measured as the angles between these planes and the MSP in the axial view (Extended Data Fig. 1-6).

TDA was measured on a cross-sectional basis, where each cerebral hemisphere was independently and evenly cut into 60 slices perpendicular to the antero-posterior axis. For each slice, a bounding box covering the slice surface was generated with three edges parallel to the x, y, and z axes. The width of each bounding box was measured as the sectional hemispheric width at the corresponding brain slice. The sectional hemispheric perimeter was measured as the average length of the hemispheric pial surface at two ends of each slice. The profile of both width and perimeter measures for the contiguous slices represents the variation of the cerebral hemisphere external morphology along the antero-posterior direction. The difference of the measures between the left and right cerebral hemispheres reflected the bilateral variation of the cerebral surface (Extended Data Fig. 1-7).

Positional differences between the two cerebral hemispheres were computed at each vertex to quantify relative displacements along the left-right, antero-posterior and ventro-dorsal axes (Extended Data Fig. 1-8). The left-right positional asymmetry (Asym_LR_) was computed as the distance of a vertex on the left hemispheric surface to the MSP subtracted from that of its corresponding vertex on the right hemispheric surface. The antero-posterior positional asymmetry (Asym_AP_) and dorsal-ventral positional asymmetry (Asym_DV_) were calculated as the projections of the displacement vector between the left and right corresponding vertices along the anterior-posterior and dorso-ventral axes. In particular, the vertex-wise interhemispheric correspondences were obtained by nonlinearly registering both hemispheric surfaces to the same symmetrical reference based on local folding patterns^68^ (Extended Data Fig. 1-3). These vertex-wise SPA measures were smoothed on the tessellated surfaces using a Gaussian kernel with the full width half maximum (FWHM) of 20 mm to increase the signal-to-noise ratio and to reduce the impact of mis-registration.

All the BT measures, except the ones of frontal and occipital bending, were divided by the scaling factor obtained in the FSL linear spatial normalization so as to obtain measures in the native space. All the BT profiles were measured using programs written in-house in Matlab (R2019b). All the procedure for MRI processing and BT measurements were implemented on the LONI pipeline system for high-performance parallel computing (**Error! Hyperlink reference not valid**., ^70^. To exclude possible outliers in terms of BT measures, we excluded the subjects with at least one lobar measure exceeded 3 standard deviations from the mean in each individual cohort.

### Statistical analysis for BT and effects of age, sex, handedness and TIV

To determine population-level, average patterns of BT, one-sample t-tests were applied to individual and pooled data sets to examine the null hypothesis that the mean of each BT measure differs from zero. To characterize the complex (linear and nonlinear), lifespan age trajectories of BT using the pooled sample, linear regression models were constructed with a step-down model selection procedure testing for cubic, quadratic and linear age effects, for each lobar profile, at each brain slice for sectional TDA, and at each cortical surface vertex for SPA. Such method has been commonly used in previous studies of complex brain structural trajectories in neurodevelopment^71^ and aging^47, 72^. The full model for a BT measure *T* is

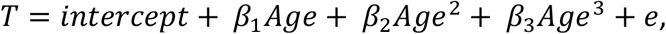

where *e* is the residual error, and the intercept and *β* terms are the fixed effects. If the cubic age effect was not significant, the cubic term was removed and we stepped down to the quadratic model and so on. The analyses were repeated controlling for sex, handedness and total Intracranial volume (TIV) as covariates. MRI acquisition sites and scanners were not included because of their pronounced associations with age in the pooled sample (P < 5e-324). To investigate effects of sex, handedness, sex-by-handedness interaction and TIV, we repeated the linear regressions with these terms as the fixed effects and adjusting for cubic, quadratic and linear age effects. Sex and handedness differences in the prevalence of BT features were examined using Chi-squared two-sample tests. We corrected for multiple comparisons for BT measures at different scales separately. For lobar and sectional measures, Bonferroni correction was applied to set a significance threshold adjusting for the number of profiles (P < 0.05/6 lobar measures and P < 0.05/60 *×* 2 sectional measures). Vertex-wise statistical results were corrected across the cortical surface using the random field theory (RFT) method^73^, which adapts to spatial correlations of the surface data. The critical threshold was set to RFT-corrected P < 0.05/3 to further correct for the 3 positional asymmetry dimensions. All linear regression analyses were conducted using our Neuroimaging PheWAS system, which is a cloud-computing platform for big-data, brainwide imaging association studies (for details, see http://phewas.loni.usc.edu/phewas/ and ^74^).

### Genomic Data and Processing

This study employed genomic data from the ABCD, PING, PNC and UKB datasets. ABCD participants were genotyped using the Affymetrix NIDA Smokescreen array, consisting of 733,293 SNP markers. For PING participants, genotyping was performed on the Illumina Human660W-Quad BeadChip with 550,000 SNPs. Of the initial 1,445 PNC participants recruited for brain imaging, 657 were genotyped on the Illumina HumanHap 610 array; 399 on the Illumina HumanHap 550 array; 281 on the Illumina Human Omni Express array and 108 on the Affymetrix Axiom array. Prior to imputation, the ABCD, PING and PNC datasets were filtered so as to exclude samples with low individual call rate < 90%, sex discrepancy, outlying heterozygosity (> mean*±*3SD), non-European ancestry (determined by population stratification) and/or being related to another sample (PI_HAT > 0.2), and to remove variants with high missingness (>10%), low minor allele frequency (MAF < 0.05) and/or strong deviation from Hardy-Weinberg equilibrium (P < 1e-5). Then the filtered genotyping datasets were phased with Eagle v2.4 and imputed using Minimac4 for the Haplotype Reference Consortium (HRC) r1.1 reference Panel via the Michigan Imputation Server^75^ (https://imputationserver.sph.umich.edu/).

UK Biobank participants were genotyped using the Affymetrix UK BiLEVE Axiom array (on an initial ∼50,000 participants) and the Affymetrix UK Biobank Axiom array (on the remaining ∼450,000 participants). The two arrays are very similar with over 95% common marker content. In total, about 800,000 markers were genotyped for each participant. The UK Biobank team first imputed the genotyping data using the Haplotype Reference Consortium (HRC) reference panel and then imputed the SNPs not in the HRC panel using a combined UK10K + 1000 Genomes panel. The imputation process produced a dataset (V3, released in March 2018) with >92 million autosomal SNPs. Detailed information and documentation on the genotyping, imputation and QC are available at (http://www.ukbiobank.ac.uk/scientists-3/genetic-data/). Post-imputation sample and variant quality control (QC) was performed to delete subjects missing more than 10% of total imputed genotypes and without valid BT measures, and to remove multiallelic variants and variants with an imputation information score < 0.7, missingness rate > 10%, MAF < 1% and/or HWE P < 1e-7. The processed datasets consisted of 1,399 ABCD samples with 7,016,694 variants, 350 PING participants with 7,383,086 variants, 422 PNC samples with 7,224,238 variants and 18,206 UKB participants with 11,195,104 variants. The pre- and post-imputation QC described above was performed using the PLINK2.0 software^76^ (https://www.cog-genomics.org/plink/2.0/).

### Heritability estimation

We estimated the narrow-sense (pedigree-based) heritability of the BT features using the twin data of the ABCD and HCP cohorts. For ABCD samples, zygosity was determined using the probability of identity-by-descent (IBD) extracted from the genome-wide genotyping data of 2,652 subjects with brain MRI. According to the protocol introduced in the ABCD data release note - NDA 2.0.1 Genetics, we identified full siblings or dizygotic twins with a threshold of IBD > 0.4 and monozygotic twins with a threshold of IBD > 0.8 rather at 0.5 and 1, considering the existence of inherent noise in the genotyping data caused by the identify-by-state estimations. 120 monozygotic twin pairs and 379 full sibling or dizygotic twin pairs were identified in the ABCD participants. In HCP S900 samples, twin statuses were self-reported. However, in the later S1200 release, zygosity was updated based on genotyping data available from blood and saliva samples of some HCP subjects. 36 twin pairs who self-reported as dizygotic twins were found to be genetically monozygotic. Taking this change into account, the S900 subjects with brain MRI included 179 twin pairs (114 MZ with 99 siblings and 6 half siblings and 65 DZ with 55 siblings and 9 half siblings), 273 siblings, 10 half siblings and 87 unpaired individuals. Pedigree-based heritability was estimated using the variance component analysis method as implemented in SOLAR-Eclipse^77^(v8.4.2, http://solar-eclipse-genetics.org). Each BT measure was entered into a linear mixed-effects model as a dependent variable. The model included fixed effects of age, age^2^, sex, handedness and TIV, and a random effect of genetic similarity that was determined by pedigrees. Genetic similarity is coded as 1 for MZ twins who share 100% of their DNA sequence, and as 0.5 for DZ twins and siblings who share on average 50%, and as 0 for unrelated individuals. Heritability statistics were corrected for multiple comparisons using Bonferroni correction for lobar and sectional measures adjusting for the number of profiles (P < 0.05/6 lobar measures and P < 0.05/60 *×* 2 sectional measures) and using false discovery rate (FDR) for vertex-wise results (FDR < 0.05/3 across the cortical surface and adjusting for the 3 positional asymmetry dimensions).

We also estimated SNP-based heritability of the BT features using the genome-based restricted maximum likelihood (GREML) analysis implemented in GCTA^78^(v1.93.2beta, https://cnsgenomics.com/software/gcta/#Overview) with the QCed genotyping dataset from UKB and using the LD-score regression (LDSC)^79^ (https://github.com/bulik/ldsc) with the GWAS summary statistics for UKB and the meta-GWAS summary statistics. The analysis for SNP-based heritability was not applied to the ABCD, PNC and PING cohorts because of their insufficient sample sizes in the filtered genomic datasets, which would result in nonsensical estimates, i.e., 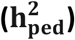values will be out of the 0-100% range and/or with very large variance (SE >> 10%).

### Genome-wide association analysis

Genome-wide association study (GWAS) analyses on different BT measures were conducted for the ABCD, PING, PNC, UKB samples using generalized linear regression adjusting for age, age^2^, sex, handedness, TIV, the first ten genetic principal components and MRI acquisition parameters if needed, as implemented in the PLINK2.0. The cubic age term was not included in the model as there was no cubic age effect in the individual samples. In addition to the lobar and sectional measures, we computed mean Asym_LR_, Asym_AP_ and Asym_DV_ within 74 brain regions defined by the Destrieux atlas ^80^ as the traits for SPA, in order to overcome the computational challenge to implement vertex-wise GWASs. We then used METAL^33^ (v2020-05-05, https://genome.sph.umich.edu/wiki/METAL) to perform meta-analyses using the z-scores method, based on p-values, sample size and direction of effect, with genomic control correction. We discarded variants that existed in only one individual GWAS, providing overall P-values on 7,393,766 high-quality genotyped and imputed autosomal SNPs. Significant genetic associations were first identified using the commonly used genome-wide significance threshold P < 5e-08, which accounts for the number of SNPs tested in modern GWAS and the correlation structure between SNPs in European ancestry populations^81^. For a more conservative discovery, we further adjusted the threshold with a Bonferroni factor that accounts for the number of BT measures tested (6 lobar measures, 60 *×* 2 sectional TDA measures, and 74 *×* 3 regional mean SPA measures), giving a threshold of P < 5e-8/348 = 1.44e-10. The clumping function in PLINK using an LD *r*^2^ cut off of 0.2 and a 500 kb sliding window was implemented to identify independent leading SNPs in each LD block. Association lookups were performed for all leading SNPs via the NHGRI-EBI catalog of published GWASs^34^ (http://www.ebi.ac.uk/gwas/) to check whether these SNPs had been previously associated with any traits.

### Gene-based association analysis

Gene-based association analysis was conducted using MAGMA ^35^ (v1.07, https://ctg.cncr.nl/software/magma). The gene-based statistics were derived using the summary statistics from each meta-analysis. Genetic variants were assigned to genes based on their position according to the NCBI 37.3 build, with a gene window of 5 kb upstream and downstream of transcript end sites for each gene, resulting a total of 18,359 protein coding genes. The European panel of the 1000 Genomes data^82^ (phase 1, release 3) was used as a reference panel to account for linkage disequilibrium. A significance threshold for gene-based associations was calculated using the Bonferroni method to correct for multiple testing across the genes (P < 0.05/18359 = 2.72e-6), and a further Bonferroni adjustment was also applied for the number of BT measures tested (P < 0.05/18359/348 = 7.83-9). Association lookups was performed in the NHGRI-EBI GWAS catalog again to explore previously reported associations of the genes identified here.

### Gene-set enrichment analysis

MAGMA gene-set enrichment analysis^83, 84^ was performed on the genome-wide, gene-based statistics to explore implicated biological pathways, examining 7,563 Gene Ontology ‘biological process’ sets from the Molecular Signatures Database^85^ (MSigDB, v7.2, https://www.gsea-msigdb.org/gsea/msigdb). Briefly, for each BT profile, the gene-wise association statistics computed in the above gene-based association analysis were quantified as Z-scores using a probit transformation mapping stronger associations onto higher Z-scores. Then, for each gene set, a linear regression model was implemented on a gene-level data matrix of gene-set indicator variables (coded 1 for genes in that gene set and 0 otherwise) to test whether the mean association of genes in the gene set is greater than that of genes not in the gene set. Significant gene-set enrichments were assessed at a significance threshold adjusting for multiple testing across the gene sets (P < 0.05/7563 = 6.61e-6), and further adjusting for the number of BT measures tested (P < 0.05/7563/348 = 1.90e-9).

### Gene-property analysis

We also preformed MAGMA gene-property analysis^83, 84^ on the genome-wide, gene-based statistics to assess relationships between the gene-BT associations in our data and the gene expression levels at different brain developmental stages from the BrainSpan data^36^ (https://www.brainspan.org). The BrainSpan dataset comprised brain tissue gene expression values at 31 age stages, from 8 postconceptional weeks (pcw) to 40 years. The gene-property analysis is essentially the same model as the above competitive gene-set analysis, but using continuous gene-set predictor variables (gene expression values at each age stage here) rather than binary indicators. This approach tested for the degree to which the genetic association of a gene changes as the value for the tested variable increases^84^. Significance of gene property effects were determined controlling for multiple testing across the age groups (P < 0.05/31 = 0.0016), and further controlling for the number of BT measures tested (P < 0.05/31/348 = 4.63e-6).

### Genetic correlation analysis

We estimated genetic correlations between the BT features studied here and 9 complex traits that were widely related to brain asymmetry previously. We fetched GWAS summary statistics of recent large-scale studies on Educational Attainment^86^ (N = 1,131,881), Intelligence^87^ (N = 269,867), attention deficit hyperactivity disorder^88^ (N=55,374), autism spectrum disorder^89^ (N = 46,350), Schizophrenia^90^ (N = 306,011), Bipolar Disorder^91^ (N = 51,710), Major Depression^92^ (N = 480,359), Neuroticism^93^ (N = 449,484) and Alzheimer’s Disease^94^ (N = 455,258), from https://www.thessgac.org/data, https://www.med.unc.edu/pgc/ and https://ctg.cncr.nl/software/summary_statistics. These summary statistics were correlated with the meta-GWAS summary statistics for BT measures obtained in this study using LDSC^79^. Significant genetic correlations were tested using a significant threshold correcting for the 9 traits and 348 BT measures tested here (P < 0.05/9/348 = 1.60e-5).

### Phenome scan analyses

Phenome scan analyses were conducted for the BT measures derived from the UKB dataset using the Phenome Scan Analysis Tool^32^ (PHESANT, https://github.com/MRCIEU/PHESANT) so as to search for other related variables in addition to age, sex, handedness and genetic factors. 5,791 phenomic variables of sociodemographics, lifestyle, environment, cognitive functions, health and medical information, and medical imaging markers were extracted from the UKB database. PHESANT was specifically designed for performing comprehensive phenome scans across the complex UKB data fields, using an automated, rule-based method to determine how to test variables of different types. A detailed description of PHESANT’s automated rule-based method is given in ^32^. PHESANT estimates the bivariate association of each BT measure with each phenotypic variable in the dataset using linear, logistic, ordered logistic, and multinominal logistic regression for continuous, binary, ordered categorical, and unordered categorical data types respectively. Prior to testing, an inverse normal rank transform is applied to variables of the continuous data type, to ensure they are normally distributed. All analyses were adjusted for covariates as in the GWAS analyses described beforehand. Phenome-wide significance was determined using a critical threshold adjusting for the 5,791 phenotypes and the 348 BT measures (P < 0.05/6334/348 = 2.27e-8).

## Data Availability

The data used in this work were obtained from 6 publicly available datasets: ABCD, HCP, ICBM, PING, PNC and UKB (application number 25641). We also used 9 publicly available GWAS summary statistics from several GWAS databases. All these data resources were described and cited in the Methods section. The genomic and phenomic analyses summary statistics are given in Supplementary Table 8-22.

## Code Availability

This study used openly available software and codes as described in the text, including FSL v5.0 (https://fsl.fmrib.ox.ac.uk/fsl/fslwiki/), FreeSurfer v6.0 (https://surfer.nmr.mgh.harvard.edu), LONI pipeline (http://pipeline.loni.usc.edu), Neuroimaging PheWAS (http://phewas.loni.usc.edu/phewas/), PLINK v2.0 (https://www.cog-genomics.org/plink/2.0/), Michigan Imputation Server (https://imputationserver.sph.umich.edu/), SOLAR-Eclipse v8.4.2 (http://solar-eclipse-genetics.org), GCTA v1.93.2beta (https://cnsgenomics.com/software/gcta/#Overview), LDSC (https://github.com/bulik/ldsc), METAL v2020-05-05 (https://genome.sph.umich.edu/wiki/METAL), MAGMA v1.07 (https://ctg.cncr.nl/software/magma) and PHESANT (https://github.com/MRCIEU/PHESANT). The in-house Matlab programs for BT measurements are available from the authors upon request.

## Author contributions

L.Z.: Conceptualization, methodology, analysis, bioinformatics, visualization, original draft writing, review & editing. W.M.: Methodology, analysis, review & editing. Y.S.: bioinformatics, review & editing. R.P.C.: bioinformatics, review & editing. T.W.A: Conceptualization, direction, funding acquisition, supervision, review & editing.

## Acknowledgements

This work was supported by the Big Data for Discovery Science (BDDS) (NIH Grant No. U54EB020406), the Laboratory of Neuro Imaging Resource (LONIR) (NIH Grant No. P41EB015922), and the Genetic Influences on Human Neuroanatomical Shapes (NIH Grant No. R01MH094343).

Data used in this work was obtained from the Adolescent Brain Cognitive Development (ABCD), the Human Connectome Project (HCP), the International Consortium for Brain Mapping (ICBM), the Pediatric Imaging, Neurocognition, and Genetics (PING), the Philadelphia Neurodevelopmental Cohort (PNC) and the UK Biobank Resource (UKB). ABCD is supported by the National Institutes of Health and additional federal partners under award numbers U01DA041048, U01DA050989, U01DA051016, U01DA041022, U01DA051018, U01DA051037, U01DA050987, U01DA041174, U01DA041106, U01DA041117, U01DA041028, U01DA041134, U01DA050988, U01DA051039, U01DA041156, U01DA041025, U01DA041120, U01DA051038, U01DA041148, U01DA041093, U01DA041089, U24DA041123, and U24DA041147. A full list of supporters is available at https://abcdstudy.org/federal-partners.html. A listing of participating sites and a complete listing of the study investigators can be found at https://abcdstudy.org/consortium_members/. HCP (WU-Minn Consortium, Principal Investigators: David Van Essen and Kamil Ugurbil; 1U54MH091657) is funded by the 16 NIH Institutes and Centers that support the NIH Blueprint for Neuroscience Research; and by the McDonnell Center for Systems Neuroscience at Washington University. ICBM (Principal Investigator: John Mazziotta, MD, PhD) was supported by the National Institute of Biomedical Imaging and BioEngineering. ICBM data are disseminated by the Laboratory of Neuro Imaging at the University of Southern California. ICBM is the result of efforts of co-investigators from UCLA, Montreal Neurologic Institute, University of Texas at San Antonio, and the Institute of Medicine, Juelich/Heinrich Heine University – Germany. PING was supported by the National Institute on Drug Abuse and the National Institute of Child Health and Human Development (RC2DA029475, R01HD061414). PNC was supported by MH089983 and MH089924. This research was conducted, using the UK Biobank Resource under approved project 25641 (principal applicant: L.Z.). The investigators within ABCD, HCP, ICBM, PING, PNC and UKB provided data but did not participate in analysis or writing of this article.

Particular thanks to Dr. Lily Xiang for the assistance with building the workflow for computing complex BT measures, and Dr. Xiang-Zhen Kong for sharing the manuscript on brain skew study^16^ .

---

**Extended Data Figure 1:**
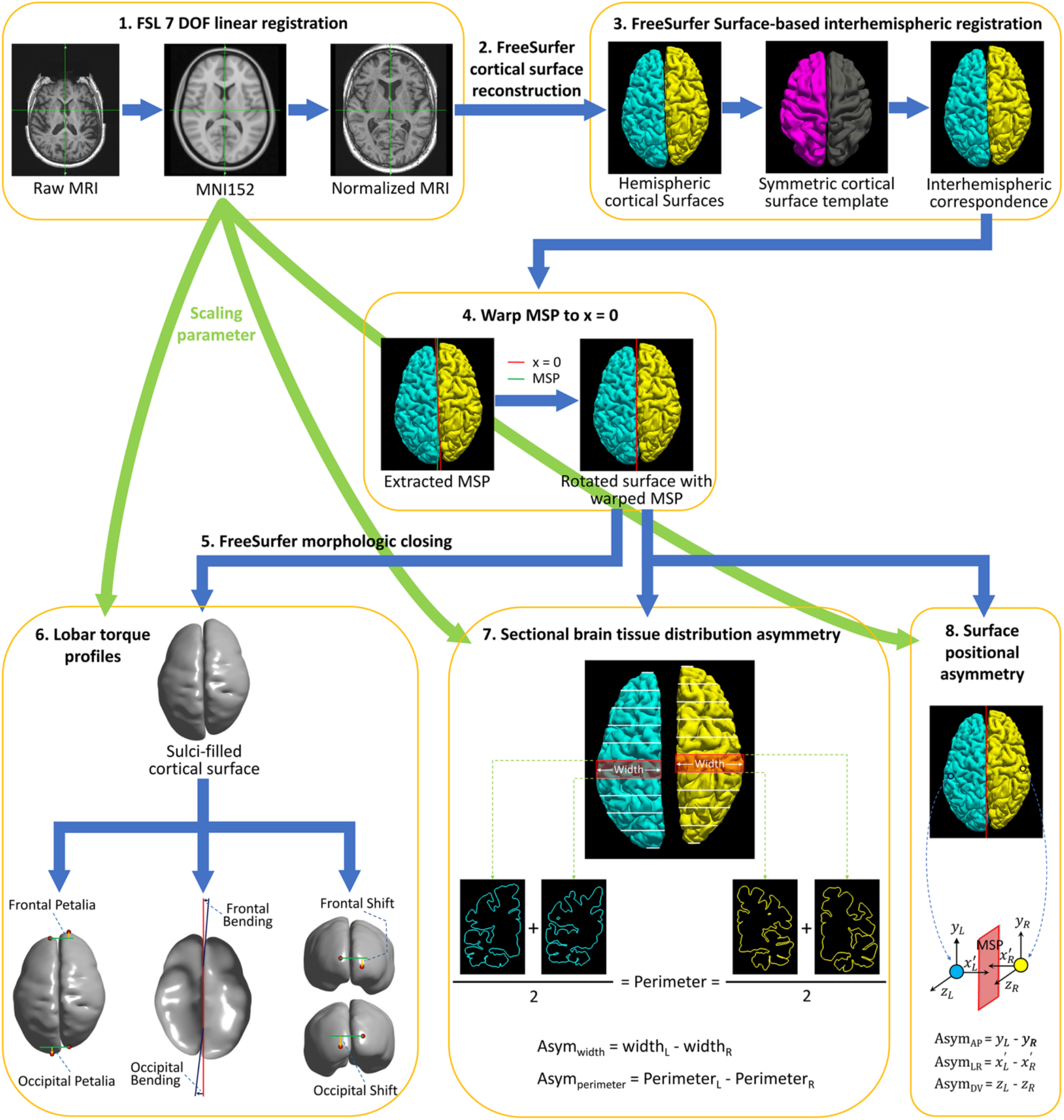
Workflow for computing complex brain torque profiles. Step 1: FSL preprocessing for normalizing the native brain volume to the standard MNI152 template using a 7 degrees of freedom (DOF) transformation. Step 2: FreeSurfer processing on the normalized brain volume to reconstruct cortical hemispheric surfaces. Step 3: FreeSurfer surface-based interhemispheric registration to establish vertex-wise interhemispheric correspondence. Step 4: Correcting for mis-registration of the brain mid-sagittal plane (MSP) by rotating the brain surface with an 3D angle between MSP and the plane x = 0. MSP was extracted as the plane best fitting the two hemispheric medial surfaces. 5. FreeSurfer morphologic closing to fill sulci and smooth the cortical surface. Step 6: Computing petalia and shift as the respective displacements of the left and right frontal/occipital extreme points along the antero-posterior and dorso-ventral axes respectively. Estimating bending as the angles between the planes best fitting the interhemispheric fissure in the frontal/occipital regions and MSP. Step 7: Measuring left-right asymmetries in hemispheric width (Asym_width_) and perimeter (Asym_perimeter_) in 60 contiguous coronal brain slices. Sectional hemispheric width and perimeter was measured as the bounding box width and the average pial surface length at two ends of each brain slice respectively. Step 8: Computing interhemispheric surface positional asymmetries along the left-right axis (Asym_LR_) as vertex-wise differences in the distances to MSP on the x-axis, and asymmetries along the antero-posterior (Asym_AP_) and dorsal-ventral axis (Asym_DV_) as the vertex-wise relative displacements on the y- and z-axis respectively.

## Legends of Supplementary Tables and Figures

**Supplementary Table 1:** Sample characteristics in the individual and pooled datasets.

**Supplementary Table 2:** Population-level average petalia, bending and shift in the individual and pooled datasets.

**Supplementary Table 3:** Prevalence of different configurations of frontal/occipital petalia, bending and shift in the studied sample.

**Supplementary Table 4:** Sex and handedness differences in prevalence of brain torque configurations.

**Supplementary Table 5:** Sex and handedness differences in prevalence of single brain torque profiles.

**Supplementary Table 6:** SNP-based Heritability of lobar brain torque features estimated using the LDSC and GCTA-GREML methods. LDSC was applied to the GWAS summary statistics for the UK biobank (UKB) cohort and the meta-GWAS summary statistics. GCTA-GREML was applied to the UKB genomic data.

**Supplementary Table 7:** List of anatomical parcellations defined by the FreeSurfer Destrieux atlas.

**Supplementary Table 8:** Genetic associations identified in meta-GWAS at the significance level of P < 5e-8. Independent lead SNPs are highlighted in bold. Associations survived in further adjustment at P < 5e-8/348 = 1.44e-10 are highlighted in red.

**Supplementary Table 9:** GWAS Catalog results for traits previously associated with identified genes (lead SNPs) influencing brain torque (BT).

**Supplementary Table 10:** GWAS Catalog results for previously reported associations of the identified genes (lead SNPs) influencing brain torque (BT).

**Supplementary Table 11:** Significant gene-based associations (P < 0.05/18359 genes = 2.72e-06) identified using MAGMA gene-based analysis.

**Supplementary Table 12:** GWAS Catalog results for traits previously associated with the genes identified using MAGMA gene-based analysis.

**Supplementary Table 13:** GWAS Catalog results for previously reported associations of the identified genes identified using MAGMA gene-based analysis.

**Supplementary Table 14:** Significant gene-set associations (P < 0.05/7563 gene sets = 6.61e-6) identified using MAGMA gene-set analysis.

**Supplementary Table 15:** Relations (P < 0.05/31 = 0.0016) between gene-based associations with brain torque features and higher gene expression levels in the human brain at particular ages.

**Supplementary Table 16:** Trends of genetic correlations between brain torque features and other traits (P < 0.05).

**Supplementary Table 17:** Phenome-wide associations for frontal/occipital petalia, bending and shift (P<0.05/6334=7.89e-6). Associations with P < 0.05/6334/348 = 2.27e-8 are highlighted in green.

**Supplementary Table 18:** Phenome-wide associations for hemispheric width asymmetries (P<0.05/6334=7.89e-6). Associations with P < 0.05/6334/348 = 2.27e-8 are highlighted in green.

**Supplementary Table 19:** Phenome-wide associations for hemispheric perimeter asymmetries (P<0.05/6334=7.89e-6). Associations with P < 0.05/6334/348 = 2.27e-8 are highlighted in green.

**Supplementary Table 20:** Phenome-wide associations for surface positional asymmetries along the antero-posterior axis (Asym_AP) (P<0.05/6334=7.89e-6). Associations with P < 0.05/6334/348 = 2.27e-8 are highlighted in green.

**Supplementary Table 21:** Phenome-wide associations for surface positional asymmetries along the left-right axis (Asym_LR) (P<0.05/6334=7.89e-6). Associations with P < 0.05/6334/348 = 2.27e-8 are highlighted in green.

**Supplementary Table 22:** Phenome-wide associations for surface positional asymmetries along the dorsal-ventral axis (Asym_DV) (P<0.05/6334=7.89e-6). Associations with P < 0.05/6334/348

= 2.27e-8 are highlighted in green.

**Supplementary Table 23:** List of self-reported neurological and psychiatric disorders that were excluded in the current study.

**Supplementary Figure 1:** Plots of occipital (x-axis) and frontal (y-axis) petalia, bending and shift with 95% confidence ellipses in the pooled sample. Population prevalence of each brain torque configuration is annotated in the plots.

**Supplementary Figure 2-A:** Population-level average sectional width asymmetry in the individual and pooled datasets. Significant sectional asymmetries (P < 0.05/120 and asymmetry/(left+right) > 2%) are marked with black dots.

**Supplementary Figure 2-B:** Population-level average sectional perimeter asymmetry in the individual and pooled datasets. Significant sectional asymmetries (P < 0.05/120 and asymmetry/(left+right) > 2%) are marked with black dots.

**Supplementary Figure 3-A:** Population-level average surface positional asymmetry along the left-right axis in the individual and pooled datasets. Color bars represent T statistics (thresholded at random field theory corrected P < 0.05/3). Red-yellow shows leftward and rightward displacements of the left hemisphere relative to the right respectively.

**Supplementary Figure 3-B:** Population-level average surface positional asymmetry along the antero-posterior axis in the individual and pooled datasets. Color bars represent T statistics (thresholded at random field theory corrected P < 0.05/3). Red-yellow and blue-cyan show forward and backward displacements of the left hemisphere relative to the right respectively.

**Supplementary Figure 3-C:** Population-level average surface positional asymmetry along the dorso-ventral axis in the individual and pooled datasets. Color bars represent T statistics (thresholded at random field theory corrected P < 0.05/3). Red-yellow and blue-cyan show upward and downward displacements of the left hemisphere relative to the right respectively.

**Supplementary Figure 4:** SNP-based heritability of sectional asymmetries of hemispheric width (A) and perimeter (B) estimated using the LDSC and GCTA-GREML methods. LDSC was applied to the GWAS summary statistics for the UK biobank (UKB) cohort and the meta-GWAS summary statistics. GCTA-GREML was applied to the UKB genomic data. Blue dots represent the heritability estimates; error bars show the standard errors; symbol ‘x’ presents nonsensical estimates of h^2^ < 0.

**Supplementary Figure 5:** SNP-based heritability of regional mean surface positional asymmetries along the left-right (Asym_LR_), antero-posterior (Asym_AP_) and dorsal-ventral (Asym_DV_) axes estimated using the LDSC (A) and GCTA-GREML (B) methods. LDSC was applied to the GWAS summary statistics for the UK biobank (UKB) cohort and the meta-GWAS summary statistics. GCTA-GREML was applied to the UKB genomic data.

**Supplementary Figure 6:** Manhattan plots of meta-GWAS summary statistics for the brain torque profiles showing significant genetic associations. Red and blue lines represent the significance levels of P < 5e-8 and P < 5e-8/348 = 1.44e-10 respectively. Independent lead variants are annotated with their SNP IDs and the closest genes. LocusZoom plots (purple diamond symbols indicate lead SNPs) are attached to show regional associations. For the abbreviations of brain regions, see Supplementary Table S7.

**Supplementary Figure 7:** Q-Q plots of meta-GWAS summary statistics for the brain torque profiles showing significant genetic associations. For the abbreviations of brain regions, see Supplementary Table S7.

**Supplementary Figure 8:** Overview of MAGMA gene-property analysis results with respect to the BrainSpan age-specific gene expression data. Ribbons in the connectogram plot illustrate associations (P < 0.05/31) of gene expression levels at 31 neurodevelopmental stages (left) with the genetic associations with brain torque (BT) profiles (right). Different colors represent different age stages or BT profiles. Ribbons are colored with the colors of linked BT profiles. Ribbon thickness represent the -log(P) value of the corresponding association. AsymWidth = Width Asymmetry, AsymPerimeter = Perimeter Asymmetry, AsymLR = asymmetry along left-right axis, AsymAP = asymmetry along Antero-Posterior axis, AsymDV = asymmetry along Dorso-Ventral axis. For the abbreviations of brain regions, see Supplementary Table S7.

**Supplementary Figure 9:** Relation between gene-based associations with brain torque features and higher gene expression levels in the brain at particular ages, using BrainSpan data from 31 age groups. Asterisks represent P < 0.05/31 = 0.0016. For the abbreviations of brain regions, see Supplementary Table S7.

**Supplementary Figure 10:** Genetic correlations between brain torque (BT) features and other traits. Only BT features showing at least one trend (P < 0.05) of genetic correlation (highlighted with asterisks) are included here.

**Supplementary Figure 11:** Sex and handedness differences in fluid intelligence (verbal-numerical reasoning) score in the UK Biobank cohort.

